# Heterodimerization of Endolysin Isoforms During Bacterial Infection by Staphylococcal Phage φ2638A

**DOI:** 10.1101/2024.01.16.575832

**Authors:** Léa V. Zinsli, Anna M. Sobieraj, Patrick Ernst, Susanne Meile, Samuel Kilcher, Cedric Iseli, Anja Keller, Birgit Dreier, Peer R. E. Mittl, Andreas Plückthun, Martin J. Loessner, Mathias Schmelcher, Matthew Dunne

**Affiliations:** Institute of Food Nutrition and Health, ETH Zurich, Switzerland; Department of Biochemistry, University of Zurich, Switzerland

## Abstract

Bacteriophage endolysins targeting Gram-positive bacteria typically feature a modular architecture of one or more enzymatically active domains (EADs) and cell wall binding domains (CBDs). Several endolysins also feature internal translational start sites (iTSSs) that produce short variant (SV) isoforms alongside the full-length (FL) endolysin. While the lytic activity of endolysins and their isoforms has been extensively studied as exogenous agents, the purpose behind producing the SV isoform during the phage infection cycle remains to be explored. In this study, we used staphylococcal phage φ2638A as a model to determine the interplay between its full-length endolysin, Ply2638A, and its SV isoform during phage infection. X-ray crystallography structures and AlphaFold-generated models enabled elucidation of individual functions of the M23 endopeptidase, central amidase, and SH3b domains of Ply2638A. Production of the SV isoform (amidase and SH3b) was confirmed during phage infection and shown to form a heterodimer complex with Ply2638A via inter-amidase domain interactions. Using genetically engineered phage variants, we show that production of both isoforms provides an advantage during phage infection as phages producing only one isoform presented impaired lytic activity, which could be partly restored through recombinant protein complementation of the missing isoform. Importantly, when applied as an antimicrobial protein against *Staphylococcus aureus* in culture, the activity of Ply2638A remained constant regardless of SV isoform complementation. Drawing from our findings, we propose that SV isoform production provides its biological advantage upon endolysin entry to the periplasmic space to ensure optimal peptidoglycan degradation prior to cell wall lysis and progeny phage release.

## Introduction

Endolysins are bacteriophage-encoded peptidoglycan (PG) hydrolases produced during the end of the bacteriophage lytic cycle to degrade the bacterial cell wall and enable the release of progeny virions. Given the global emergence and spread of antimicrobial resistance (AMR) (WHO 2020) there is significant interest in developing endolysins as precision antimicrobial agents for treatment of bacterial infections and chronic conditions, where antibiotics lack effectiveness or are actively contraindicated due to antibiotic stewardship initiatives. Recombinant endolysins effectively target Gram-positive bacterial pathogens, such as *Staphylococcus aureus,* due to their ability to directly access and degrade the exposed PG layers (Fischetti 2010). This distinct mode of action compared to antibiotics also renders endolysins highly effective against antibiotic resistant pathogens, such as methicillin-resistant *S. aureus* (MRSA), which caused more than 120,000 deaths attributable to antimicrobial resistance in 2019 (Murray et al. 2022), as well as dormant bacteria and biofilms (Gutierrez et al. 2014, Olsen et al. 2018).

Endolysins targeting Gram-positive bacteria typically feature a modular architecture (Schmelcher et al. 2012) consisting of one or more enzymatically active domains (EAD), cell wall binding domains (CBDs) used for substrate recognition, and flexible linkers of variable length connecting these domains (Diaz et al. 1990, Schmelcher et al. 2012). The EADs have evolved to cleave specific PG bonds found within their target bacterial cell walls, and contain activities such as, (i) *N*-acetyl-β-D-muramidases, (ii) lytic transglycosylases and (iii) *N*-acetyl-β-D-glucosaminidases, which all cleave bonds within the sugar backbone of the PG; (iv) *N*-acetylmuramoyl-L-alanine amidases, which cleave the amide bond between sugar and peptide moieties; and (v) endopeptidases, which cleave different peptide bonds within the stem peptide or interpeptide bridge connecting the sugar backbones of the PG (Schmelcher et al. 2012). The combination of EADs with the ability to cleave specific peptidoglycan structures, along with genus-, species-, or serovar-specific CBDs, is among the key advantages driving the interest in developing endolysins as precision antimicrobials capable of targeting pathogenic species while leaving commensal microbiomes unaffected (Fowler et al. 2020, Son et al. 2020, Danis-Wlodarczyk et al. 2021). There is substantial structural and functional diversity among endolysins (Rahman et al. 2021). Comprehending the interplay between these diverse domains will not only facilitate our efforts to engineer endolysin-based bacteriolytic agents, but also helps unravel the molecular intricacies of how endolysins naturally function during the final stages of phage lysis and their entry into the periplasm, which in Gram-positive bacteria describes the space between the cell membrane and the peptidoglycan.

Endolysins targeting Gram-positive bacteria frequently possess in-frame, internal translational start sites (iTSSs), resulting in the expression of two endolysin isoforms (Pinto et al. 2022). These iTSS-harboring endolysins can be categorized into two types: Type I, which have an iTSS between two EADs, leading to the production of an enzymatic short variant (SV) isoform, consisting of a single EAD and C-terminal CBD, along with the full-length (FL) endolysin (Catalao et al. 2011, Abaev et al. 2013); and Type II, which have an iTSS between the last EAD and CBD, resulting in the co-expression of the CBD alone alongside the FL endolysin (Proenca et al. 2015, Dunne et al. 2016, Zhou et al. 2020). Multimers of varying FL and SV stoichiometries have been observed for Type II iTSS endolysins (Proenca et al. 2015, Dunne et al. 2016), where the SV isoforms form complexes with FL endolysins, such as heterotetramers of CTP1L (2FL:2SV) (Dunne et al. 2016) and LysIME-EF1 (1FL:3SV) (Zhou et al. 2020), representing the final form of the endolysin that has maximum lytic activity. However, little is known about the role of the SV isoform produced by Type I iTSS endolysins and their potential interactions or complex formation with the FL isoform (Pinto et al. 2022).

Staphylococcal endolysins typically have a three-domain architecture consisting of an N-terminal endopeptidase, a central *N*-acetylmuramoyl-L-alanine amidase, and a C-terminal CBD of SH3b functional homology (Schmelcher et al. 2015). The endopeptidase is often a cysteine/histidine-dependent aminohydrolase/peptidase (CHAP) domain, as reported for a variety of well-studied staphylococcal endolysins from phages GH15 (Gu et al. 2011), Twort (Loessner et al. 1998), and phi11 (Navarre et al. 1999). The endolysin Ply2638A from *Staphylococcus pseudintermedius* phage φ2638A uses an N-terminal M23 endopeptidase instead of a CHAP domain (Abaev et al. 2013), but interestingly, this domain has evolved to target the same PG cleavage site as the CHAP domains of the other staphylococcal endolysins mentioned above (Schmelcher et al. 2015). Ply2638A also has a Type I iTSS, which leads to co-expression of an SV isoform consisting of the amidase and CBD alongside the FL isoform (Abaev et al. 2013). To the best of our knowledge, iTSSs have not been identified in conventional CHAP-containing staphylococcal endolysins. The Ply2638A native isoform combination (termed within as Ply_WT_) has shown high *in vitro* activity against *S. aureus* and the ability to rescue mice from MRSA-induced septicemia (Schmelcher et al. 2015). Using the Ply2638A composition as a scaffold, more effective three-domain chimeric enzybiotics have subsequently been engineered for therapeutic applications, including Staphefekt SA.100 and XZ.700, with the latter shown to be effective at reducing bacterial numbers in different *S. aureus*-induced skin infection models (Eichenseher et al. 2022, Pallesen et al. 2023) as well as for removing biofilms from titanium discs mimicking prosthetic joint infections (Kuiper et al. 2021).

While Ply2638A and other endolysins targeting different bacterial pathogens have shown their potential as enzybiotic applied exogenously, substantial knowledge gaps remain in our understanding of the interplay between endolysin domains and their isoforms, especially during phage lysis and PG degradation. For instance, the inherent benefit of Type I iTSS endolysins producing both a truncated and active SV isoform has remained elusive. In this study, Ply2638A served as a model Type I iTSS endolysin, enabling the structural characterization of the FL endolysin and a functional investigation of its interplay with the SV isoform. We aimed to understand why, when expressed together by the parental phage φ2638A, they exhibited greater bacteriolytic activity, with our investigation unveiling a surprising interaction between the two isoforms through their central amidase domains that may have implications for other Type I iTSS endolysins.

## Results

### Optimal phage fitness requires expression of both Ply2638A isoforms

After confirming the *in vivo* functionality of the *ply2638A* iTSS (Figure 1B), we aimed to explore the role of both isoforms during phage infection and lysis. *S. pseudointermedius* 2854 cultures were infected with φ2638A WT, φ2638A *ply*_FL_, or φ2638A *ply*_SV_ (no hemagglutinin (HA)-tag) (Figure 1A) and the reduction in optical density (OD_600_) of the bacterial culture was measured over eight hours using turbidity reduction assays (TRAs) (Figure 1C). The wildtype phage exhibited the most effective infection dynamics, reaching a maximum OD_600_ of 0.48 within 2.3 hours followed by a sharp and sustained reduction in optical density until a slight regrowth was observed near the end of the experiment (∼6.5 hours). In contrast, both engineered phages, φ2638A *ply*_FL_ and φ2638A *ply*_SV_, exhibited a noticeable decrease in bacteriolytic activity compared to wildtype. In both cases, the bacterial culture was able to grow to higher maximum OD_600_ values of 0.71 and 0.73 within 3.1 and 3.3 hours, respectively, with the delay in lysis leading to the turbidity of both infections plateauing at higher optical densities. The reduced bacteriolytic activity of both engineered phages was also observed at different ratios of phage to bacteria (∼4 x 10^7^ CFU/mL), defined as the multiplicity of infection (MOI) ranging from 0.001 to 1.0. Differences in activity became negligible when phage quantities exceeded ten-fold the bacterial count (MOI >10) (Supplementary Figure S1). The difference in bacteriolytic activity between these phages was also assessed using time-kill assays (TKAs) by measuring absolute colony-forming unit (CFU) survival over six hours (Supplementary Figure S2). Surprisingly, lower phage concentrations (MOIs of 0.1 or 1.0) resulted in minimal bacterial killing by all three phages despite revealing clear differences in bacteriolytic activity when measuring OD_600_ changes using TRAs. To address this, a higher MOI of 10 was tested, even though all three phages demonstrated similar activity profiles by TRA at this higher phage titer. Indeed, all three phages displayed the same 4-log reduction in CFU levels within the first hour, with no significant variation in bacterial counts over the remaining six hours. Post-plating phage infection or endolysin activity could potentially further obscure any differentiation in bacteriolytic activity between the three phages using TKAs. During phage production, no differences were observed in the final yields of either engineered phage in comparison to the wildtype. This suggests that the absence of either isoform does not markedly affect progeny release, indicating that any discernible differences in activity are likely to be subtle. Consequently, we proceeded with using TRAs and a standardized MOI of 0.1 to further explore phage activity and the role of the two endolysin isoforms.

**Figure 1.**
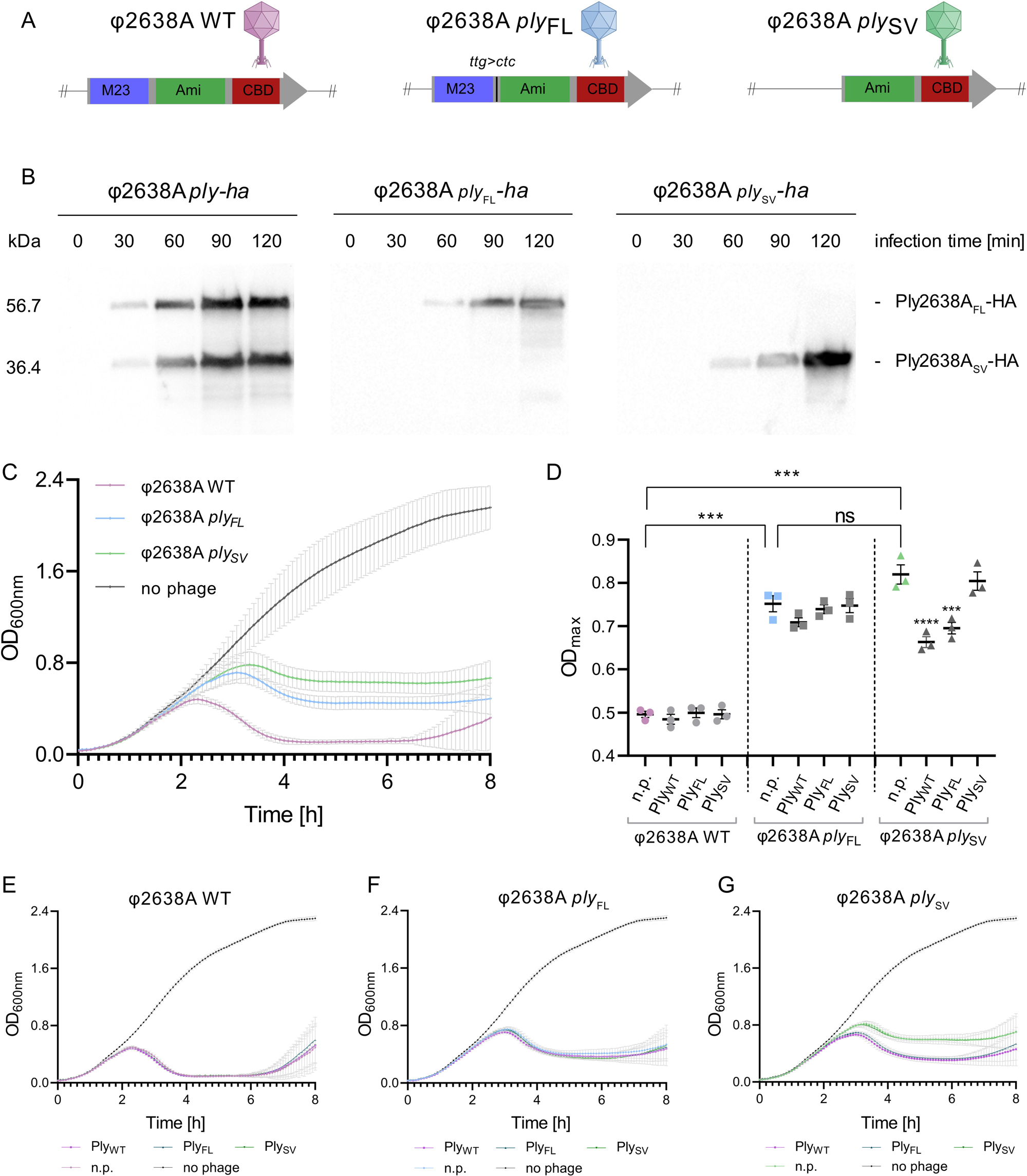
Ply2638A short variant isoform is required for maximum bacteriolytic activity during phage infection. **(A)** Schematic overview of the endolysin genes of wildtype and engineered φ2638A phages. M23, M23 peptidase; Ami, amidase; CBD, SH3b cell wall binding domain; *ttg>ctc*, codon modification that silences the iTSS to produce only FL endolysin. **(B)** Western blot time-course using an anti-HA monoclonal antibody to monitor hemagglutinin (HA)-tagged endolysin expression after phage infection of *S. pseudointermedius* 2854 cultures. **(C)** Bacteriolytic activity of wildtype and engineered phages against *S. pseudointermedius* 2854 was determined by 8-hour turbidity reduction assays (TRAs) at OD_600nm_. Phages were added at an MOI of 0.1 (10^7^ PFU/mL to 10^8^ CFU/mL). **(D-G)** Phage infections were supplemented with 10 nM of recombinant Ply_WT_, Ply_FL_, or PLy_SV_ or no protein (n.p.) as a control at the start of infection, with bacteriolytic activity measured using TRAs as described above. **(D)** The maximum optical density reached during individual infections is reported with or without endolysin complementation. Phage-only infected cultures (n.p.) are shown in color and were compared against each other using unpaired t-tests (ns, no significance; ***, p-value = 0.0002). For each phage, the three endolysin-complemented conditions were compared against the phage-only control using a one-way ANOVA. Only significant differences are indicated with asterisks (***: p-value = 0.0006; ****: < 0.0001). For panels C to E, all experiments were performed as biological triplicates with technical triplicates and shown as mean ± standard deviation.

We next sought to investigate if bacteriolytic activity of the engineered phages could be restored by supplementing phage infection with purified, recombinant Ply_WT_ (containing both isoforms), or Ply_FL_ and Ply_SV_ alone at three different concentrations (10 nM, 100 nM, and 1 µM) providing additional exogenous enzymatic activity against the cell wall. When the wildtype phage was complemented with 10 or 100 nM of any endolysin variant, no discernible differences in turbidity reduction were observed when compared to phage infection alone (n.p., no protein control) (Figure 1E). Similar trends were observed for endolysin complementation of φ2638A *ply_FL_*, where no increase in bacteriolytic activity was evident compared to the phage-only conditions (Figure 1F). In contrast, when phage φ2638A *ply_SV_* was complemented with either Ply_WT_ or Ply_FL_, turbidity reduction was more efficient compared to the phage alone or after supplementation with Ply_SV_ with the growth curve exhibiting characteristics of φ2638A *ply_FL_* infection (Figure 1G). These observations were further supported by decreases in OD_600_ max values (Figure 1D), which dropped to 0.66 (p-value < 0.0001) and 0.69 (p-value 0.0006) when complemented with 10 nM Ply_WT_ or Ply_FL_, respectively, compared to the phage alone. A ten-fold increase in endolysin concentration (100 nM) produced similar effects, with no change for the wildtype or φ2638A *ply_FL_* but an increase again in the bacteriolytic activity of φ2638A *ply_SV_* complemented with either Ply_WT_ or Ply_FL_ (Supplementary Figure S3). Interestingly, supplementation of any phage with 1 µM Ply_SV_ negatively affected the bacteriolytic activity of all three phages, with minimal variation observed for 1 µM Ply_FL_ or Ply_WT_ complementation, suggesting that Ply_SV_, at higher (atypical) concentrations than expected during phage infection, may interfere with phage infection and/or the ability to effectively lyse bacterial cells (Supplementary Figure S3).

### Ply_FL_ and Ply_SV_ form an inter-amidase domain heterodimer

Structural investigations of endolysins harbouring a Type II iTSS, producing SV isoforms consisting of the CBD alone, have unveiled distinct multimeric complexes between the two isoforms, for instance, clostridial endolysin CTP1L forms a heterotetrameric complex (2 FL and 2 SV isoforms) (Dunne et al. 2016) (PDB ID: 5A6S) and enterococcal endolysins LysIME-EF1 (Zhou et al. 2020) (PDB ID: 6IST) and Lys170 (Xu et al. 2021) (PDB ID: 7D55) assemble as heteropentameric complexes, featuring one FL and four SV isoforms. In contrast, the formation of heteromeric complexes by endolysins featuring a type I iTSSs, producing SV isoforms of active EAD-CBD fusions, has not been explored, and no evidence of multimerization has been observed either in solution or in crystal structures. Hence, our discovery of isoform co-elution during the size exclusion chromatography (SEC) of Ply_WT_ for the phage complementation assays (Supplementary Figure S4), warranted further investigation.

Size-exclusion chromatography combined with multi-angle light scattering (SEC-MALS) was used to investigate potential complex formation between the two isoforms (Figure 2A). When analyzed separately, each isoform exhibited a single peak close to their monomeric masses: Ply_FL_ at 56.9 ± 0.3 kDa (expected mass, 55.5 kDa) and Ply_SV_ at 40.8 ± 0.3 kDa (expected mass, 35.3 kDa) (Figure 2A). However, upon combining an excess of Ply_SV_ with Ply_FL_ (w/w ratio of 3:1), a higher molecular species, suggestive of a heterodimer, was identified at a mass of 97 ± 1.2 kDa. This indicated that all Ply_FL_ molecules were engaged with a single Ply_SV_, while the unbound Ply_SV_ retained its monomeric state, with a calculated molecular weight of 42.6 ±0.4 kDa. Complex formation did not occur at Ply_FL_ to Ply_SV_ ratios of 1:1 or 3:1 (Supplementary Figure S5), implying that an excess of Ply_SV_ was necessary for complex formation under the tested conditions. Furthermore, no homodimer formation was observed for either of the isoforms tested alone or in combination.

**Figure 2.**
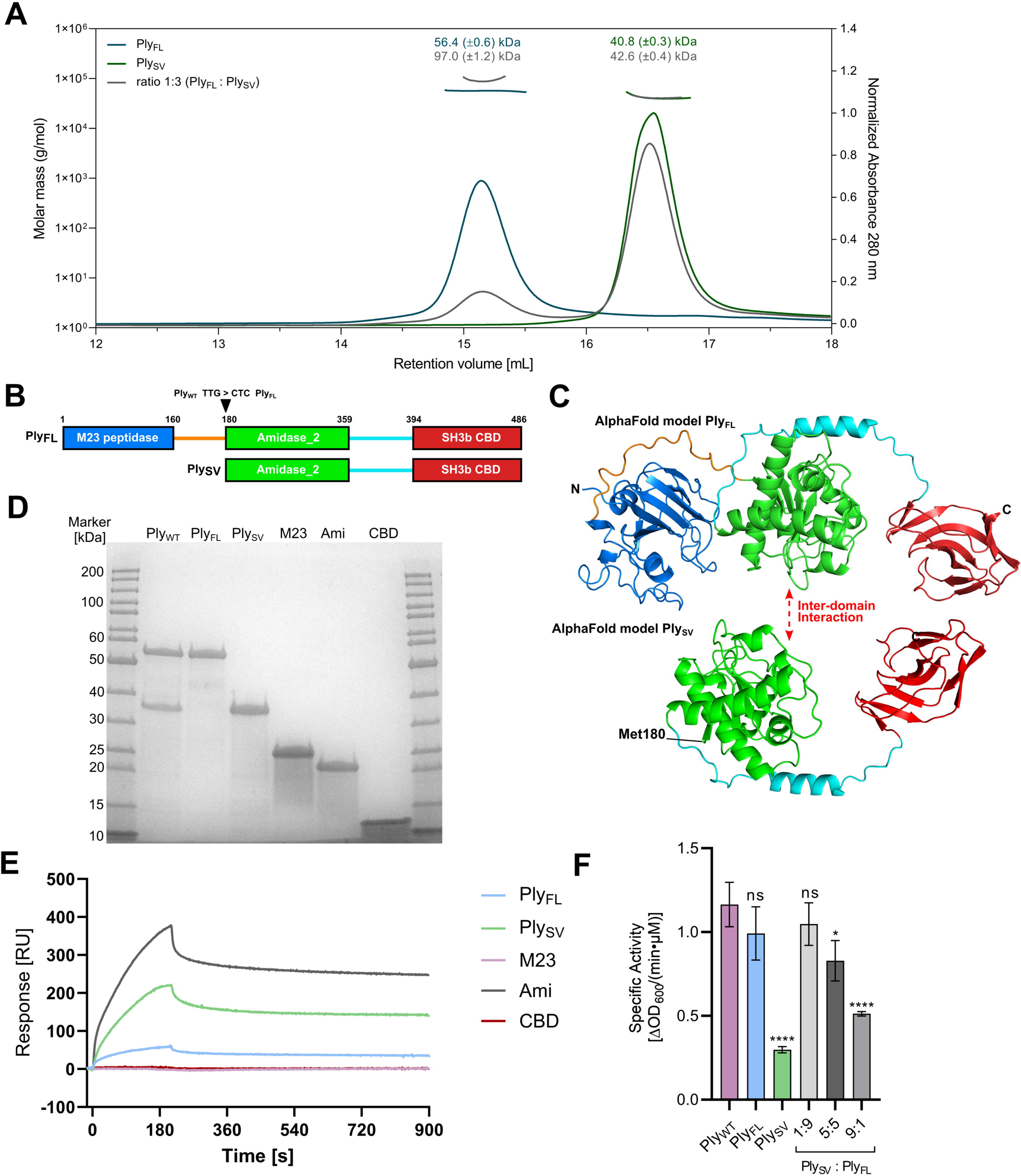
The SV isoform forms an inter-amidase domain interaction with the full-length Ply2638A. **(A)** SEC-MALS analysis of the Ply2638A heterodimer. The oligomeric state represented by Ply_FL_ (blue), Ply_SV_ (green) and a 1:3 (w/w) mixture of Ply_FL_ and Ply_SV_ (grey) at a concentration of 1 mg/mL. The curves shown represent the absorption at 280 nm (peaks; right *y*-axis). The determined mass of the Ply_FL_, Ply_SV_ and the complex are shown by the data points on top of the corresponding protein peaks (left *y*-axis). **(B)** Schematic overview of the Ply2638A and the SV isoform with residues sitting at domain and linker boundaries highlighted. The arrow indicates the SV isoform translational start site (TTG) that is silent mutated to CTC for production of Ply_FL_ alone. **(C)** AlphaFold 2.0 (Jumper et al. 2021) generated models of Ply_FL_ and Ply_SV_ colored as shown in panel B **(D)** SDS-PAGE analysis of each construct produced and purified for SPR and SEC-MALS analysis. **(E)** SPR sensorgrams of the analytes Ply_FL_ (light blue), Ply_SV_ (light green) and single domains M23 (purple), Ami (dark grey) and CBD (red) interacting with the ligand Ply_FL_ immobilized on the chip surface. Analyte concentration was 2.5 µM for all five constructs. **(F)** Specific activity of Ply_WT_ (purple), Ply_FL_ (blue), Ply_SV_ (green) and different ratios of Ply_FL_ and Ply_SV_ (grey) determined by turbidity reduction assays using *S. aureus* Cowan cells. Error bars represent standard deviations from the experiments that were performed in biological triplicates with technical triplicates each. A one-way ANOVA was performed to compare the specific activity of Ply_WT_ to the other proteins and ratios (ns: nonsignificant; *: < 0.05; ****: < 0.0001).

To investigate how these proteins were interacting, and to identify the domains involved in complex formation, binding experiments were conducted using surface plasmon resonance (SPR). Ply_FL_ was immobilized as the ligand and recombinant M23 peptidase, central amidase, SH3b CBD, as well as the Ply_FL_ and Ply_SV_ isoforms were assessed as analytes (Figure 2E; Supplementary Figure S7). Dose- and construct-dependent interactions were observed for Ply_FL_, Ply_SV_, and the amidase domain alone, with the strongest signals observed for the latter. Conversely, the CBD and M23 peptidase domains showed no interaction with Ply_FL_. Combining data from SEC-MALS and SPR thus strongly suggested heterodimeric complex formation of the two isoforms through inter-amidase domain interactions.

Building upon our observations with SEC-MALS, indicating an excess of Ply_SV_ being required for heterodimeric complex formation in solution, we performed TRAs to assess if combining the recombinant isoforms in different ratios (1:9, 1:1, 9:1) would lead to observable differences in bacteriolytic activity (Figure 2F). Interestingly, Ply_FL_ alone exhibited an equivalent level of activity as the native combination of both isoforms (Ply_WT_), which contrasted with previous findings where Ply_FL_ activity was significantly lower than Ply_WT_ when assessed under similar conditions when tested against *S. aureus* Newman (Abaev et al. 2013) instead of *S. aureus* Cowan cells used here. Additionally, we observed much lower activity for Ply_SV_ compared to Ply_WT_ and Ply_FL_, which again differed from previous observations where both isoforms exhibited comparable levels of activity (Abaev et al. 2013). Nevertheless, here the differences observed aligned more closely with the expectation that a full-length endolysin, featuring two EADs, would naturally exhibit higher activity than a single EAD-containing SV isoform. Given the comparable activity of Ply_FL_ and Ply_WT_, it was unsurprising that Ply_FL_, when present in a 9:1 excess, closely resembled the activity of Ply_WT_, and how with an increasing proportion of Ply_SV_ (at ratios of 1:1 and 9:1) overall activity decreased. These results mirrored our *in vivo* observations with φ2638A *ply_SV_* (Figure 1G), where phage complementation with the more active Ply_FL_ led to an overall improvement in bacteriolytic activity. Overall, while the presence of both isoforms enhances bacteriolytic activity during native phage infection, there seems to be no discernible advantage in combining Ply_FL_ with the SV isoform when applied exogenously to staphylococcal cells under the current testing conditions.

### Structural analysis of Ply2638A and its individual domains

The discovery of heterodimerization between the Ply2638A isoforms led us to assess the structural relationship of the amidase domains through X-ray crystallography. Despite multiple attempts, crystallization of the native Ply_WT_ mixture, as well as Ply_FL_ and Ply_SV_ individually, proved unsuccessful. Consequently, we shifted our focus to crystallizing the individual domains of Ply2638A. Crystals diffracting to 2.3 Å and 2.5 Å were used to determine the structures of the M23 endopeptidase (residues 1-174) (Figure 3) and SH3b (residues 393-486) domains (Figure 4). However, despite extensive testing, we were unable to obtain diffraction-quality crystals of the central amidase domain (residues 180-359). Despite this setback, AlphaFold 2.0 (Jumper et al. 2021) was used to generate models of the amidase domain (Figure 5) as well as the two Ply2638A isoforms (Figure 2C). All three models presented high per-residue confidence scores (pLDDTs) of 87.7 (Ply_FL_), 90.1 (Ply_SV_), and 96.5 (Amidase), indicating their suitability for structural assessment. AlphaFold-Multimer (Evans et al. 2022) was used to predict an amidase homodimer as well as the heterodimer of Ply_FL_ and Ply_SV_; however, all models presented poor interface pTM scores (typically below 0.2) with many predictions containing non-permissible features of a transient dimer such as intertwined loops and non-adjacent amidases domains, which prompted the omission of these models from further analysis.

#### M23 peptidase domain features a restricted substrate recognition site

The M23 peptidase of Ply2638A (Met1-Ala156) features the conserved β-sheet core structure and catalytic motif of H(x)_n_D (for Ply2638A, n=3) and HxH that is shared across this well-characterized family of zinc-dependent metallopeptidases (Małecki et al. 2021, Razew et al. 2022) (Figure 3A). Flanking the highly conserved core are four variable loops (L1-L4) that create the walls of the negatively charged binding groove whose composition determines peptidoglycan binding specificity (Małecki et al. 2021) (Figure 3 B,C). Interestingly, both Loop 1 and 4 of Ply2638A are approximately twice as long as the corresponding loops in available crystal structures of other related M23 peptidases (Figure 3 E,F). Specifically, Loop 1 in Ply2638A spans 34 residues (Asp18 to Ala51) whereas equivalent loops in the domain of structurally related bacteriocins lysostaphin (LST) (PDB ID: 4QPB and 4LXC; 17 residues) (Sabala et al. 2014) and LytM (PDB ID: 4ZYB; 13 residues) (Grabowska et al. 2015), as well as the endolysin EnpA of *Enterococcus faecalis* phage 03 (PDB ID: 6SMK; 13 residues) (Małecki et al. 2021) are notably shorter. Similarly, Loop 4 comprises 15 residues (Pro140 to Gly154), which contrasts with the shorter loops of 7, 8, and 7 residues of these related M23 peptidase domains, respectively.

**Figure 3.**
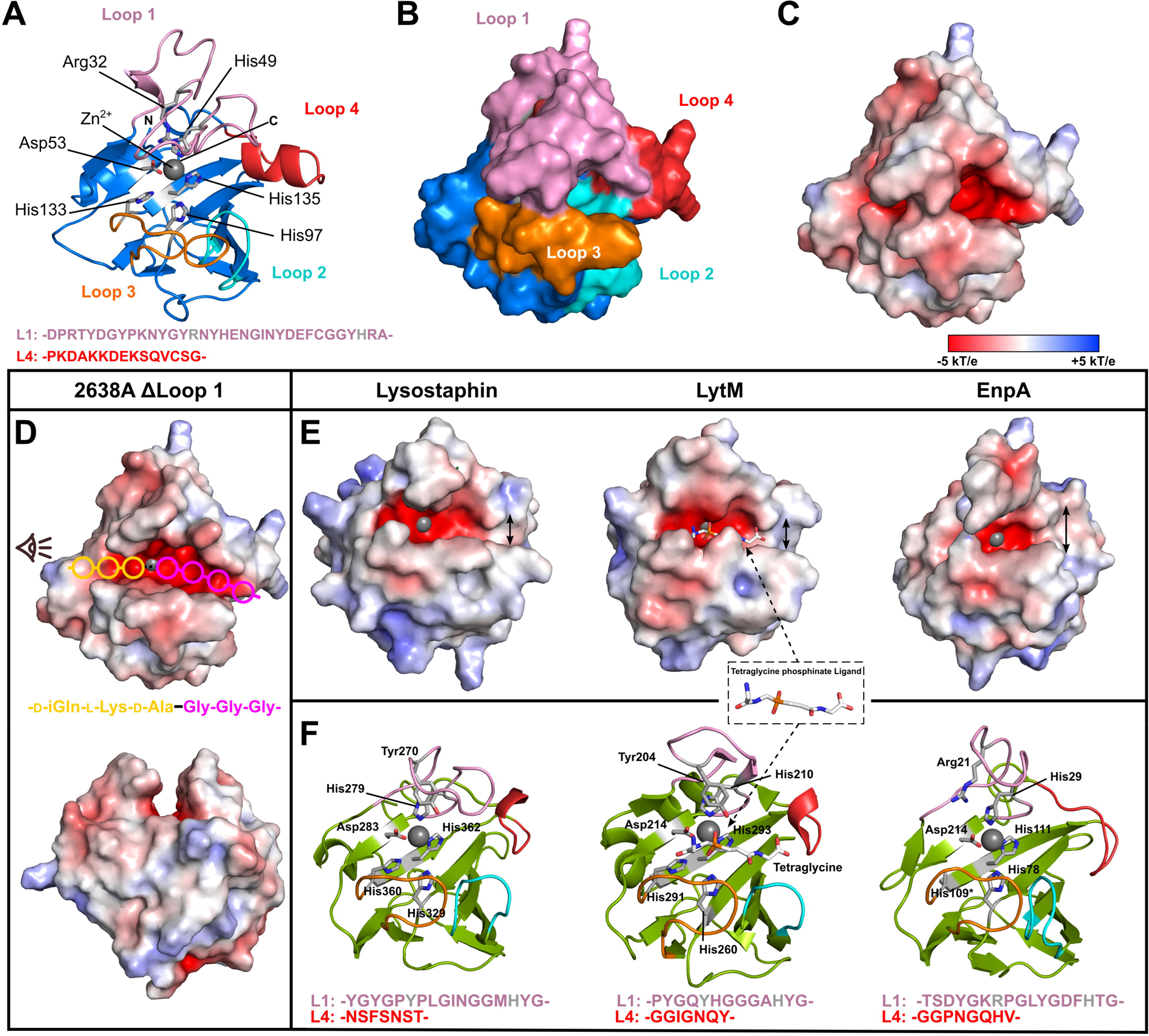
Analysis of the M23 peptidase crystal structure. Cartoon **(A)** and surface **(B)** representations of the Ply2638A M23 peptidase domain colored blue with catalytic and Zn^2+^ (grey sphere) coordinating residues colored grey and shown as sticks. The four variable loops that form the binding groove peripheral to the active site are individually colored with residues forming Loop 1 and Loop 4 indicated below. **(C)** The molecular surface of the peptidase domain colored according to its electrostatic surface potential (±5 kT/e) generated by Adaptive Poisson-Boltzmann Solver (APBS); red, negatively charged; white, neutral; and blue, positively charged regions (±5 kT/e). **(D)** Electrostatic surface potential (±5 kT/e) of the Ply2638A peptidase domain is shown with a truncated Loop 1(ΔLoop1; ΔGlu35-Tyr41) to aid visualization of the active site. Estimated positions of individual peptidoglycan residues within the active site are shown as colored circles based on the location of the tetraglycine ligand co-crystalized with LytM (PDB ID: 4ZYB; panel E) as well as previous analyses (Grabowska et al. 2015). The scissile bond between D-Ala-Gly is shown at the center as a black line. Below is the same structure oriented as indicated by the eye icon of the top image. **(E)** Electrostatic surface potential (±5 kT/e) of structurally related peptidase crystal structures of lysostaphin (PDB ID: 4LXC/4QPB), LytM (PDB ID: 4ZYB), and EnpA (PDB ID: 6SMK) (Małecki et al. 2021). Double-headed arrows highlight the widening at the end of binding groove. **(F)** Cartoon representations of the same peptidase domains colored green with active site residues colored according to panel A, and with individual Loop 1 and 4 residues indicated below.

Another uncommon feature within Loop 1 of the peptidase domain is the choice of residue used to stabilize the oxyanion intermediate of the cleavage reaction. For the majority of M23 peptidase domains this residue is a tyrosine (e.g., Tyr270 in LST and Tyr204 for LysM). In Ply2638A, however, this role is played by Arg32, a choice of amino acid akin to that found in *Helicobacter pylori* peptidase Csd3 (An et al. 2015) and EnpA. In the case of EnpA, Arg21 was additionally proposed to contribute to the stabilization of neighboring residues within the binding groove (Małecki et al. 2021). As with other M23 peptidases, Loop 1 and Loop 3 of the Ply2638A peptidase together form a deep and narrow binding groove that mediates D-alanyl-glycine endopeptidase cleavage at its center. Notably, an α-helical bulge introduced by the longer Loop 4 of Ply2638A extends the constricted and negatively charged binding groove at its terminal end (Figure 3C), which is best visualized by removing the overhanging residues of Loop 1 (Figure 3D). In contrast, LST and LysM display a less constrained configuration in this region, while EnpA widens at this location (Figure 3E). This expansion has been associated with EnpA’s ability to bind and hydrolyze a broader range of peptidoglycan structures, specifically D-Ala-Gly/Ala/Ser, which explains its bacteriolytic activity against different species, including staphylococcal and streptococcal species (Małecki et al. 2021).

Building upon prior analyses of EnpA (Małecki et al. 2021) and drawing insights from the structure of LytM co-crystallized with tetraglycine phosphinate (mimicking the ligand during cleavage) (Grabowska et al. 2015) we can estimate the placement of the scissile bond and the associated peptidoglycan residues within the active site of the Ply2638A peptidase domain (Figure 3D). Unlike LST and LytM, which serve as glycyl-glycine endopeptidases and target the comparatively simple polyglycine crossbridge, the extended binding groove of Ply2638A likely accounts for its recognition of both the stem peptide (D-Ala-L-Lys-D-Glu-L-Ala) and polyglycine crossbridge adjacent to the scissile D-Ala-Gly bond. For instance, the bulkier Loop 4 of Ply2638A extends the negatively charged groove that would best accommodate the polyglycine crossbridge. The prospect of co-crystallization with a complete peptidoglycan fragment holds promise in elucidating the intricate interplay between these loops in governing substrate specificity, particularly for Loop 1 which would naturally exhibit more flexibility than what is observed in the current static crystal structure.

#### The SH3b domain presents similar binding properties as lysostaphin

SH3b domains are one of the most common types of cell wall binding domain identified for staphylococcal endolysins (Haddad Kashani et al. 2018). The C-terminal SH3b domain of Ply2638A shares 56 % sequence identity with the LST SH3b domain and displays an almost identical structure (PDB ID: 6RJE; DALI (Holm et al. 2023) Z-score 19.2, RMSD 0.6 Å) (Mitkowski et al. 2019) consisting of nine antiparallel β-strands (β1–β10) (Figure 4A). The potent bacteriolytic activity of LST has been associated with its ability to recognize both the pentaglycine crossbridge and the peptide stem of peptidoglycan (Figure 4C) via two independent binding sites located on opposite sides of its SH3b domain (Gonzalez-Delgado et al. 2020). In addition to conservation of both binding sites, the majority of previously identified interacting residues are also present in the Ply2638A SH3b structure, suggesting a similar dual-site recognition mechanism for this endolysin (FIGURE B,D). SH3b-LST can bind (albeit at decreasing levels) peptidoglycan obtained from *S. aureus* Δ*femB* and Δ*femAB* mutants, which contain cross-bridges of three or just one glycine, respectively (Gonzalez-Delgado et al. 2020). Based on domain similarity, we hypothesized that the SH3b of Ply2638A also does not require a complete pentaglycine cross-bridge for peptidoglycan binding. To investigate this, we conducted fluorescence microscopy and quantified the relative cell binding of GFP-tagged SH3b domains from Ply2638A and LST against various *S. aureus* strains, including a Δ*femA* mutant with a single glycine within the cross-bridge (FIGURE F,G). Both GFP-tagged SH3b-2638A and SH3b-LST exhibited a significant decrease in binding, dropping to approximately 20 %, when the pentaglycine cross-bridge was reduced to three glycine residues. Furthermore, both CBDs showed minimal residual binding (<10 %) when the pentaglycine bridge was reduced to just one glycine. These results agree with previous observations for SH3b-LST. Importantly, the absence of significant differences in binding abilities between the two domains, along with their high structural similarity and preservation of the two binding sites identified for lysostaphin (including most shared residues) (Gonzalez-Delgado et al. 2020), strongly suggest that Ply2638A can also interact with the crossbridge and peptide stem of staphylococcal peptidoglycan.

**Figure 4.**
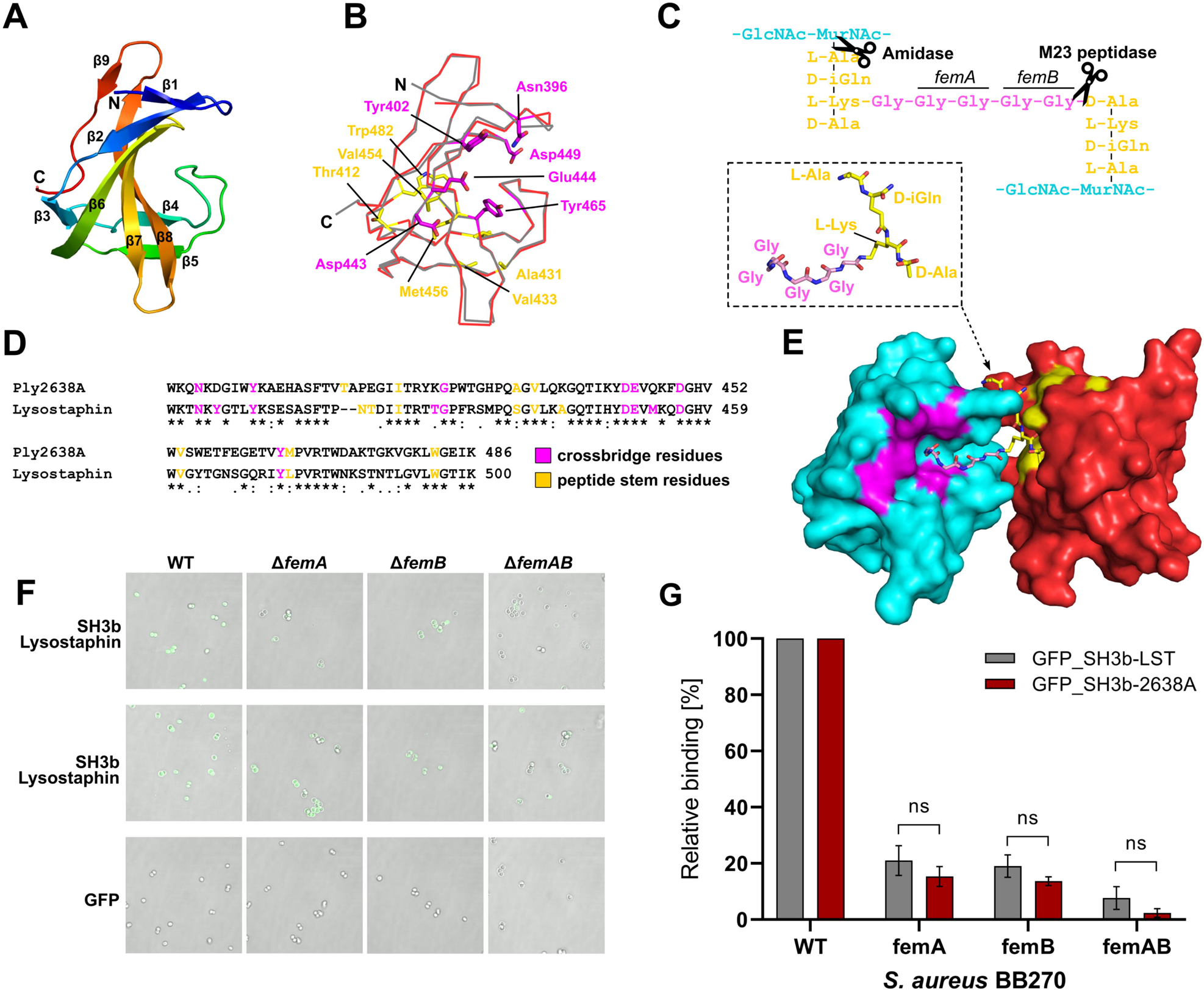
Structural and functional analysis of Ply2638A SH3b crystal structure. **(A)** Structure of Ply2638A SH3b domain (residues 393-486; PDB ID: 7AQH) colored from N-(blue) to C-terminus (red) with β-strands numbered. **(B)** Superposition of the Ply2638A (red) and lysostaphin (grey; PDB ID: 6RJE (Gonzalez-Delgado et al. 2020)) CBD with the residues previously identified for lysostaphin SH3b as interacting with the crossbridge (magenta) and the stem peptide (yellow) that are also conserved in the Ply2638A SH3b domain shown as sticks. **(C)** Schematic of the *S. aureus* peptidoglycan that is bound by the SH3b domain and cleaved (scissors) by the two enzymatic domains of Ply2638A. Enzymes *femA* and *femB* are responsible for biosynthesis of the 2^nd^ and 3^rd^ glycines and the 4^th^ and 5^th^ glycines of the crossbridge (Götz et al. 2006). **(D)** Sequence alignment of the Ply2638A and Lysostaphin SH3b domains (56 % identity) with the crossbridge and peptide stem interacting residues that are conserved between the two domains highlighted. **(E)** Surface representation of two Ply2638A SH3b domains (cyan and red) with a fragment of peptidoglycan consisting of components from the peptide stem and crossbridge (P4-G5 complex; inset) superposed using the crystal structure of lysostaphin SH3b dimers co-crystallized with P4-G5 (PDB ID: 6RJE) (Gonzalez-Delgado et al. 2020). Akin to panels B and D, crossbridge and peptide stem interacting residues are colored magenta and yellow, respectively. Representative fluorescence microscopy images **(F)** and relative (to wildtype) binding quantification **(G)** of GFP-fused SH3b domains of Ply2638A and Lysostaphin, decorating wildtype and peptidoglycan mutant strains of *S. aureus* BB270, demonstrating similar binding properties between the two SH3b domains.

#### The central amidase domain of Ply2638A

The central domain of Ply2638A (Leu180-Gly359) is a zinc-dependent, type 2 *N*-acetylmuramoyl-L-alanine amidase (IPR002502) that is responsible for cleaving the amide bond between the glycan moiety (MurNAc) and the stem peptide (L-Ala) of peptidoglycan (Figure 5 A,B). The AlphaFold model of Ply2638A amidase shares high structural similarity with available crystal structures of amidase domains of other endolysins, including those of staphylococcal phage GH15, LysGH15 (PDB ID: 4OLS; 39 % sequence identity, DALI (Holm et al. 2023) Z-score 27.2, Root Mean Square Deviation (RMSD) 1.6 Å) (Gu et al. 2014), *Bacillus anthracis* prophage Ba02, PlyL (PDB ID: 1YB0; Z-score 19.8, RMSD 2.1 Å) (Low et al. 2005), phage T7 lysozyme (PDB ID: 1LBA; Z-score 11.4, RMSD 2.6 Å) (Cheng et al. 1994), and the highly active amidase domain of *S. aureus* autolysin AtlA, AmiA (PDB ID: 4KNL; Z-score 18.2, RMSD 2.4 Å) (Büttner et al. 2014) (Figure 5D,E). The Ply2638A amidase employs the same group of zinc-binding residues, namely His206, His314, and Cys322, as the other endolysin amidases LysGH15 and PlyL (Figure 5C). Additionally, Ply2638A shares an essential catalytic glutamic acid (Glu270) with both endolysins (LysGH15, Glu282; PlyL, Glu90) as well as the AmiA autolysin (Glu324) (Büttner et al. 2014), in contrast to a tyrosine (Tyr46) as used by the T7 amidase (Cheng et al. 1994). Superposition of LysGH15 and Ply2638A revealed that the only notable compositional difference between the two amidases was an extension of ∼3 residues forming an alpha-helix within the loop between α6 and α7 of LysGH15 (Figure 5C). This loop extension was also not present in the PlyL or AmiA crystal structures, and its relevance is likely to be limited. Superimposing a muramyltetrapeptide (MtetP) resembling staphylococcal peptidoglycan, which was co-crystallized within the active site of the AmiA amidase (PDB ID: 4KNL) (Büttner et al. 2014), with the other amidases provided insights into how these different amidases recognized peptidoglycan. The three *Staphylococcus*-targeting amidases, Ply2638A, LysGH15, and AmiA, exhibit similar surface electrostatics. They feature a negatively charged binding groove and a deep active site pocket capable of accommodating MurNAc, the sugar backbone, and the stem peptide. In all three, the scissile bond is positioned in close proximity to the active site zinc ion. Despite compositional differences between MtetP and *E. coli* peptidoglycan, the *E. coli* phage T7 amidase accommodates the MtetP ligand in a similar orientation, however, with a positively charged binding groove, reflecting the enzyme’s specificity towards *E. coli* peptidoglycan (Cheng et al. 1994).

**Figure 5.**
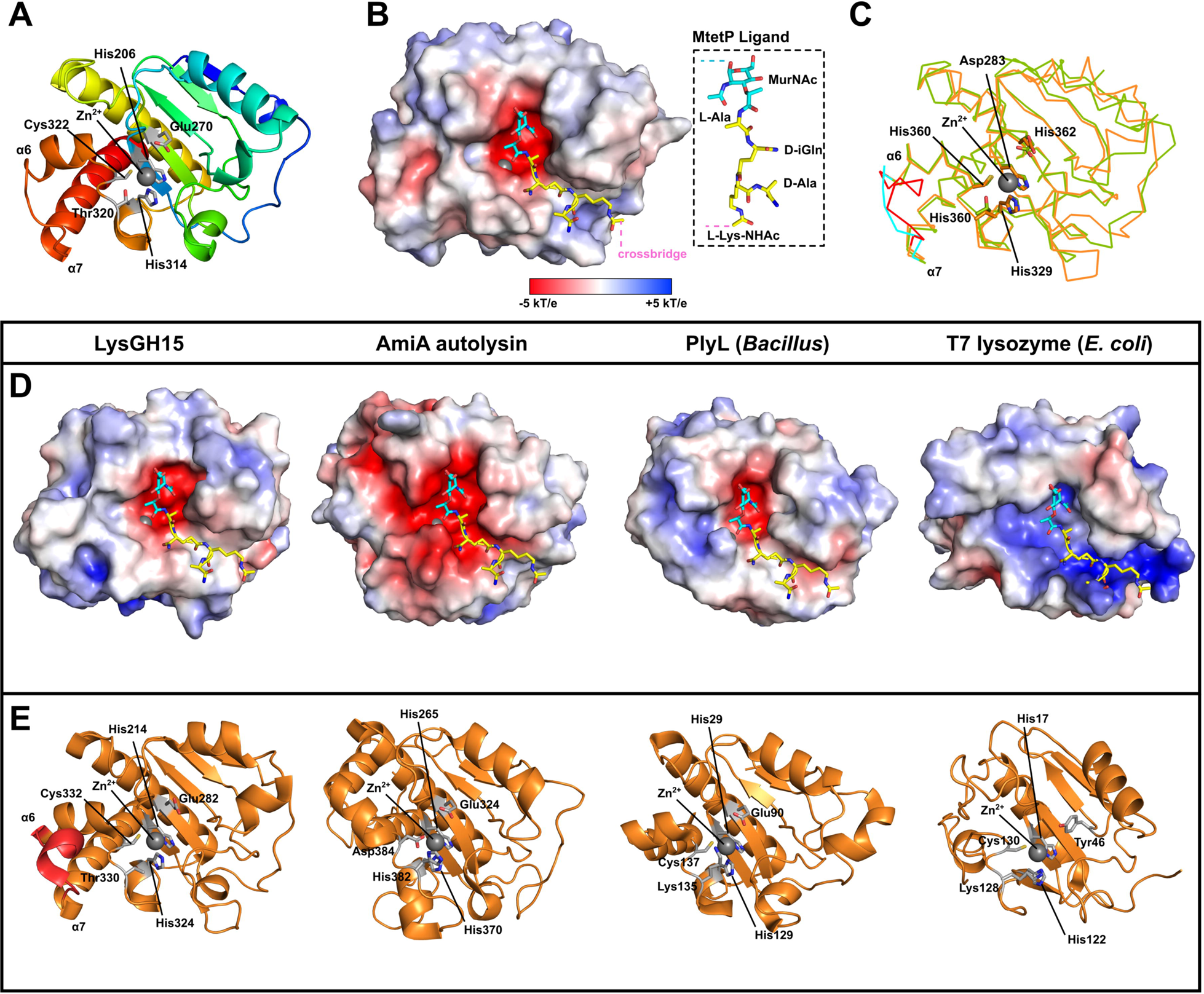
Analysis of the AlphaFold-generated model of the Ply2638A amidase domain. **(A)** Cartoon structure of Ply2638A amidase domain generated by AlphaFold 2.0 (Jumper et al. 2021) colored from N-(blue) to C-terminus (red) with catalytic and Zn^2+^ (grey sphere) coordinating residues colored white and shown as sticks. **(B)** Molecular surface of the amidase domain colored according to its electrostatic surface potential generated by APBS (±5 kT/e); red, negatively charged; white, neutral; and blue, positively charged regions. The muramyltetrapeptide (MtetP) ligand representative of *S. aureus* peptidoglycan, was modelled into the negatively charged active site by superimposing with the MtetP co-crystallized structure of the *S. aureus* autolysin, AmiA (PDB ID: 4KNL; Z-score 18.2; RMSD 2.3 Å) (Büttner et al. 2014). **(C)** Superimposing Ply2638A (green) with LysGH15 (orange; PDB ID: 4OLS) (Gu et al. 2014), which showed the highest structural similarity based on DALI analysis (Z-score 27.2; RMSD 1.6 Å) (Holm et al. 2023), reveals the same catalytic and Zn^2+^ coordinating residues, represented as sticks, in the same orientation for both structures. The loop region between α6 and α7 are colored for Ply2638A (cyan) and LysGH15 (red). **(D)** Electrostatic surface potential (±5 kT/e) of structurally similar crystal structure amidase domains from LysGH15 (Gu et al. 2014), AmiA (PDB ID: 4KNL; Z-score 18.2; RMSD 2.4 Å) (Büttner et al. 2014), the *Bacillus* prophage Ba02 endolysin, PlyL (Z-score 19.8; RMSD 2.1 Å) (Low et al. 2005), and the T7 lysozyme (PDB ID: Z-score 11.4; RMSD 2.6 Å) (Cheng et al. 1994). All structures were modeled with MtetP in the active site as performed for Ply2638A in panel B. **(E)** Cartoon representations of the same amidase domains colored orange with active site residues colored according to panel A with the α6 and α7 loop region of LysGH15 highlighted.

For most staphylococcal endolysins, which include an N-terminal CHAP domain (e.g., LysGH15) rather than an M23 peptidase (e.g., Ply2638A), the central amidase domain’s primary role has been proposed to enhance the endolysin’s affinity for target cell walls. Bacteriolytic activity, on the other hand, has been proposed as a secondary function, with the CHAP domain contributing the majority of the bacteriolytic activity for these endolysins (Son et al. 2018). In contrast, the central amidase of Ply2638A has previously demonstrated higher bacteriolytic activity than the M23 peptidase domain when either the M23 or the amidase domain was fused separately to the SH3b binding domain (Abaev et al. 2013). Here, we also observe bacteriolytic activity by Ply_SV_ (Figure 2F). Using AlphaFold 2.0, we generated high-confidence models for a representative selection of amidase domains from staphylococcal endolysins of different compositions, all of which had been previously investigated for their bacteriolytic activity and amidase functionality (Figure 6). This selection includes the CHAP-Amidase-SH3b endolysins LysK (40 % sequence similarity to Ply2638A) (Sanz-Gaitero et al. 2014), LysSA12 (41 %) (Son et al. 2018), and LysGH15 (39 %; shown in **Figure 5D**) (Gu et al. 2014), and the Amidase-SH3b endolysin LysP108 (41 %; 100% identical to LysK) (Lu et al. 2021). To the best of our knowledge, Ply2638A is the only M23-Amidase-SH3b endolysin to have its activity investigated.

**Figure 6.**
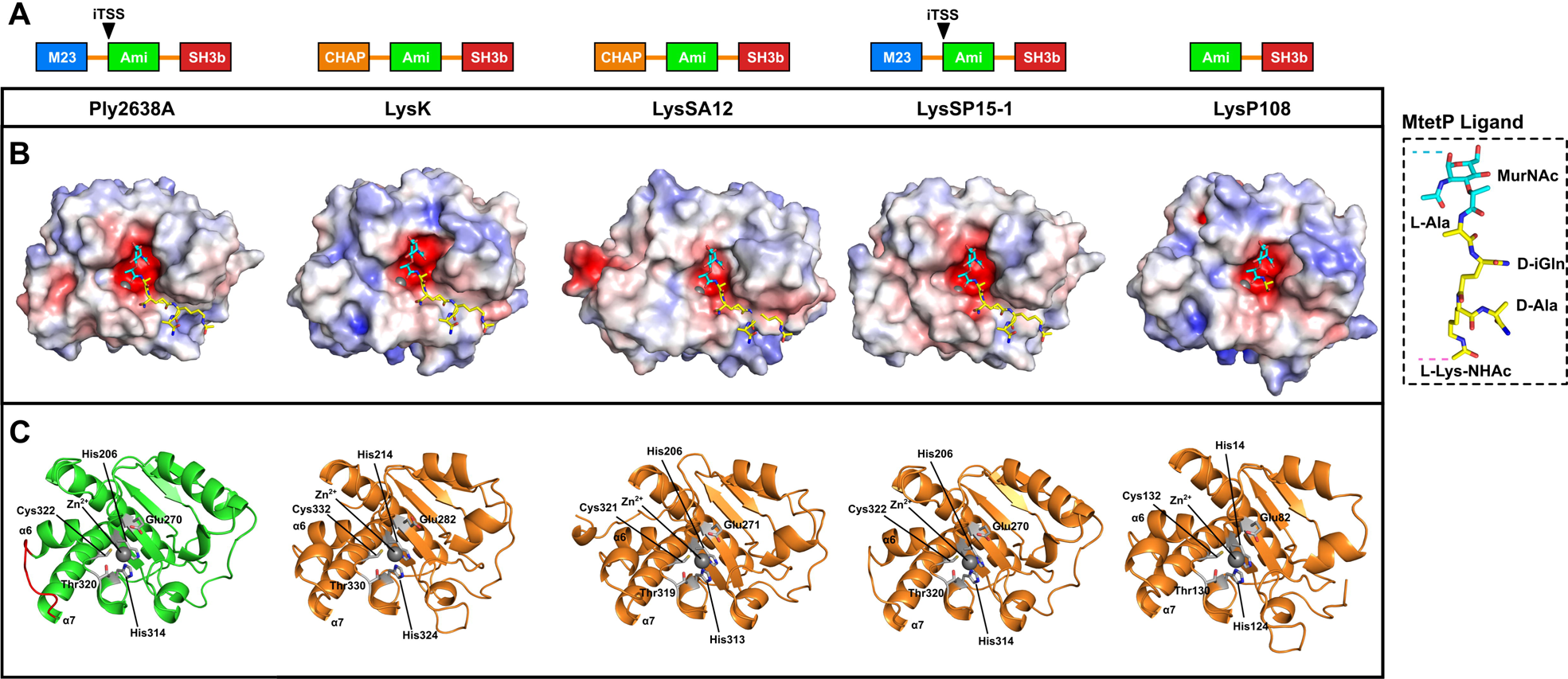
Analysis of representative amidase domains from other staphylococcal phage endolysins. AlphaFold 2.0 (Jumper et al. 2021) was used to generate high confidence models of Ply2638A (reported in Figure 5) as well as representative staphylococcal endolysins LysK (O’Flaherty et al. 2005, Becker et al. 2009), LysSA12 (Son et al. 2018), LysSP15-1 (GenBank: <u>MK075001</u>), and LysP108 (Lu et al. 2021). **(A)** LysSP15-1 features the same domain architecture as Ply2638A, whereas LysK and LysSA12 have an N-terminal Cysteine, Histidine-dependent Amidohydrolases/Peptidases (CHAP) domain instead of an M23 peptidase at the N-terminus. LysP108 features only an amidase attached to a C-terminal SH3b domain. LysSP15-1 has been predicted to also feature an internal translational start site akin to Ply2638A (Pinto et al. 2022). **(B)** Molecular surfaces of all four amidase domains colored according to their electrostatic surface potential generated by APBS (±5 kT/e); red, negatively charged; white, neutral; and blue, positively charged regions. The muramyltetrapeptide (MtetP) ligand representative of *S. aureus* peptidoglycan was modelled into the negatively charged active site by superpositioning with the MtetP co-crystallized structure of *S. aureus* autolysin, AmiA (PDB ID: 4KNL; Z-score 18.2; RMSD 2.3 Å) (Büttner et al. 2014). **(C)** Cartoon representations of the same amidase domains colored green (Ply2638A) and orange (others) with catalytic and Zn^2+^ coordinating residues colored white and shown as sticks. The α6 and α7 loop region of Ply2638A is also colored red and features an additional helical segment in LysK and LysSA12. All models presented high structural similarity to the amidase of Ply2638A with LysK, LysSA12, LysSP15-1, and LysP108 superpositioning with an RMSD (all atoms) of 0.75 Å, 0.80 Å, 0.22 Å, and 0.54 Å, respectively.

Furthermore, only a limited number of structurally analogous endolysins sourced from phage genomes were identified through BLASTp (Sayers et al. 2022). These include phages SP119-1 (GenBank AZB66744) and SPT99F3 (GenBank APD20014), both exhibiting >97 % sequence identity with Ply2638A. Consequently, we included another Ply2638A-like endolysin, LysSP15-1 (96 %; GenBank MK075001), even though its activity remains uninvestigated. Remarkably, all the amidase domains presented very high structural similarity (RMSDs between 0.2 to 1.6 Å when superimposed to the Ply2638A amidase) and contained the same active site and Zn^2+^ coordinating residues. Using the Consurf webserver (Ashkenazy et al. 2016), sequence conservation was mapped onto the Ply2638A amidase domain (Supplementary Figure S6) revealing a high degree of conservation within the negatively charged active site, contrasting with low levels of conservation across the remaining molecular surface including in close proximity to the pocket accommodating the peptide stem component of the ligand. Such surface variation might potentially explain any discrepancies in activity observed among amidase domains.

Nevertheless, establishing a connection between structural differences in these domains and their bacteriolytic activity remains challenging, particularly without direct head-to-head comparisons conducted under similar experimental conditions. As described above, Ply2638A isoform dimerization occurs via the central amidase domain. However, none of the amidase crystal structures discussed in this article, or those additionally identified as structurally similar using the DALI server, e.g., a *Bacillus subtilis* amidase (PDB ID. 3HMB; Z-score 18; RMSD 2.5 Å (Low et al. 2011)) or the *Listeria* phage PSA endolysin amidase (PDB ID: 1XOV; Z-score 19.8; RMSD 2.1 Å (Korndorfer et al. 2006)) have been characterized or reported to form homodimers. The PlyL amidase (Figure 5E) is the only exception, forming a trimer as the crystal asymmetric unit; however, this has not been described or shown to bear any functional significance.

**Table 1.** Crystallographic data statistics for M23 peptidase and SH3b domains of Ply2638A.

|  | **M23 PEPTIDASE**<br>**(AA 1-174)** | **CBD**<br>**(AA 393-486)** |
| --- | --- | --- |
| **DATA COLLECTION** | PDB ID: 6YJ1 | PDB ID: 7AQH |
| Space Group | P2<sub>1</sub> | P1 |
| Cell Dimensions |  |  |
| *a*, *b*, *c* (Å) | 33.89, 56.38, 110.49 | 62.13, 62.51, 66.171 |
| $\alpha$, $\beta$, $\gamma$ (°) | 90, 92.84, 90 | 111.01, 108.39, 90.18 |
| Wavelength (Å) | 1.00 | 1.00 |
| Resolution Range (Å) | 39.44 - 2.3 (2.38 - 2.30)<sup>‡</sup> | 44.11 - 2.49 (2.58 - 2.49)<sup>‡</sup> |
| Unique Reflections | 17,866 (1664) | 27, 918 (2454) |
| Multiplicity | 6.0 (5.7) | 2.1 (1.9) |
| Completeness (%) | 95.06 (89.46) | 90.91 (79.34) |
| mean *I*/σ*I* | 7.59 (4.10) | 4.31 (0.68) |
| Wilson B-factor | 29.1 | 51.65 |
| *R*<sub>merge</sub> | 0.1612 (0.5327) | 0.0875 (0.6781) |
| *R*<sub>meas</sub> | 0.1761 (0.585) | 0.1173 (0.9072) |
| *R*<sub>pim</sub> | 0.0696 (0.235) | 0.0775 (0.5982) |
| CC1/2 | 0.986 (0.911) | 0.992 (0.486) |
| CC* | 0.996 (0.976) | 0.998 (0.809) |
| **REFINEMENT** |  |  |
| No. Reflections used | 17,794 (1663) | 27,898 (2454) |
| Reflections for *R*<sub>free</sub> | 888 (84) | 1396 (122) |
| *R*<sub>work</sub> | 0.2414 (0.2628) | 0.2176 (0.3746) |
| *R*<sub>free</sub> | 0.3030 (0.3205) | 0.2968 (0.4182) |
| No. Atoms |  |  |
| Protein | 2713 | 6088 |
| Ligand/Ion | 2 | n/a |
| Water | 110 | 85 |
| B-factors (Å<sup>2</sup>) |  |  |
| Protein | 33.54 | 51.73 |
| Ligand/Ion | 27.64 | n/a |
| Water | 39.54 | 46.57 |
| Ramachandran Plot (%) |  |  |
| Favored | 96.12 | 93.41 |
| Outliers | 0 | 0.41 |
| R.M.S Deviations |  |  |
| Bond Lengths (Å) | 0.008 | 0.024 |
| Bond Angles (°) | 1.31 | 1.14 |

⌚ Highest resolution shell is shown in parenthesis.

## Discussion

Endolysins have emerged as a promising class of antibiotic alternatives, gaining significant attention in response to the growing antimicrobial resistance (AMR) crisis. Their alternative mechanism of action combined with their adaptability through protein engineering (Schmelcher and Loessner 2021), and species-specific activity make endolysins highly effective precision antimicrobials. Of particular interest are endolysins tailored to combat staphylococcal infections, exemplified by the clinical assessment of Exebacase and Tonabacase based on native endolysins PlySs2 (Schuch et al. 2014) and SAL-1 (Jun et al. 2014), as well as engineered endolysins SA.100 and XZ.700 based on the Ply2638A scaffold (Eichenseher et al. 2022). While the bacteriolytic properties of endolysins as enzybiotics has been well studied, the inherent role of endolysins during the latter stages of the phage lytic cycle remains largely unexplored, especially for endolysins carrying a Type I iTSS producing an enzymatically active SV isoform in addition to the FL endolysin (Catalao et al. 2011, Pinto et al. 2022).

Here, we reveal for the first time, that the Type I iTSS isoforms of Ply2638A (Abaev et al. 2013) display a transient inter-amidase interaction that mediates the assembly of heterodimeric complexes in solution. While multiple attempts to crystallize and resolve the inter-amidase interaction at atomic resolution were unsuccessful, recent breakthroughs in AI-driven prediction tools, particularly AlphaFold (Jumper et al. 2021) enabled us to generate high confidence models of both isoforms and the amidase domain alone. Unfortunately, our efforts to predict the inter-amidase interaction using AlphaFold-Multimer (Evans et al. 2022) were unsuccessful. Protein modeling has been revolutionized by such AI-driven prediction tools and excels at predicting single chain proteins and certain complexes (Humphreys et al. 2021, Gonzalez-Serrano et al. 2023, Pavlopoulos et al. 2023); however, there are still challenges at modeling more transient interfaces, like the Ply2638A inter-amidase interaction, with ongoing advancements in the field holding promise for future improvements for interface predictions (Callaway 2022, Lee et al. 2023). Consequently, in-depth atomic characterization of the inter-amidase interaction remains an area of future research.

Ply2638A inter-amidase interactions were indirectly observed via SPR analysis and SEC-MALS, and indicated the existence of a Ply_FL_:Ply_SV_ heterodimer. An interesting observation from SPR analysis was that the binding response seemed higher for constructs with fewer domains, with the amidase-only construct presenting the highest binding response. However, this finding is speculative since a steady state was not attained for the analytes, which was primarily due to high ligand (Ply_FL_) concentration immobilized on the chip surface (chosen due to the low binding capacity observed for Ply_FL_). Nevertheless, differences in binding capacities of constructs containing the interacting amidase domain can potentially be explained by the increased steric hindrance for constructs with increasing domains (i.e., apparent binding affinity of Amidase > Ply_SV_ > Ply_FL_). It is interesting to note that similar complex formations between Type I iTSS isoforms have been suggested for an *M. smegmatis* endolysin LysPollywog (also featuring a central amidase domain), however, based on inconclusive SEC analyses (Pinto et al. 2022). This implies a more widespread occurrence of this mode of interaction among endolysins; however, the requirement for forming such isoform heterodimers and the positioning of amidase domains in such complexes (i.e., whether the active sites are hidden or exposed) during the lysis process remains unknown. There is a potential scenario where, with exposed amidase active sites, the heterodimer functions in a manner similar to restriction enzyme homodimers cleaving palindromic DNA (Pingoud and Jeltsch 2001). In this hypothetical model, pairs of MurNAc-D-Ala on a cross-linked subunit of PG could potentially be cleaved simultaneously by adjacent amidase domains, facilitated by the loosening of the rigid peptidoglycan structure via the action of the single M23 peptidase domain at the N-terminus of Ply_FL_; however, additional structural data is needed to substantiate this proposition.

The requirement of Type II iTSS endolysins to maintain SV isoform production is clearly evident, as the SV isoforms of individual CBDs are essential for forming active, mature, and stable endolysin complexes (Proenca et al. 2015, Dunne et al. 2016, Zhou et al. 2020). In our study, we sought to investigate why other phages targeting Gram-positive bacteria (Pinto et al. 2022) continue to utilize a Type I iTSS, which essentially results in a less active, single EAD SV isoform that is also not required for the stability or function of the full-length isoform. The presence of the iTSS may be attributed to a gene fusion event, potentially a common trait in endolysin genes with two lytic domains, however, limited data is available to support this theory and does not explain why the iTSS would be retained when the multi-domain full-length isoform exhibits sufficiently high activity. Nevertheless, as shown here, while the wildtype and single isoform φ2638A phages exhibited similar titers during production and no discernible differences during TKA analysis measuring absolute staphylococcal killing, the wildtype phage consistently displayed superior bacteriolytic activity, as evidenced by optical density reduction via TRA analysis, in comparison to the single isoform phages. Similar results have been reported for the Mycobacteriophage Ms6, where phage mutants producing only one of two Type I iTSS isoforms were defective in the normal timing and completion of host cell lysis and also produced smaller plaques upon plating (Catalao et al. 2011).

Staphylococcal PG consists of repeating units of β-1,4-linked *N*-acetylglucosamine (GlcNAc) and *N*-acetylmuramic acid (MurNAc). These glycan strands are crosslinked by a stem peptide attached to MurNAc which is attached to another stem peptide via an interpeptide bridge (Sobral and Tomasz 2019). The PG of *S. aureus* shows a high degree of crosslinking, ranging from 74 – 92 % (Vollmer and Seligman 2010). Detailed characterization of the PG structure of *S. pseudointermedius* (the host of φ2638A) are not available leading to hypotheses based on the *S. aureus* cell wall alone. Cross-linked PG features two MurNAc-D-Ala bonds and a single L-Gly-D-Ala bond connecting the interpeptide bridge to the stem peptide, implying a twofold higher number of bonds requiring amidase cleavage compared to endopeptidase cleavage. Consequently, this may account for the necessity of Ply_SV_ co-expression, i.e., providing an additional amidase, to increase the likelihood of PG bond cleavage and facilitate more efficient bacterial cell lysis that ultimately enhances the release of progeny phages. This phenomenon was indirectly shown within this study by the higher OD_max_ observed by TRA when infecting *S. pseudointermedius* with either of the single isoform phages compared to the wildtype phage. This is attributed to a larger proportion of cells with partially degraded cell walls due to the reduced capacity to fully degrade the PG, resulting from the lack of endopeptidase activity for φ2638A *ply_SV_* or the insufficient amidase activity by φ2638A *ply_FL_*.

Although both single isoforms demonstrated the ability to induce lysis and produce phage progeny, the subtle enhancement in PG degradation represents a valuable marginal advantage for dual-isoform Type I iTSS endolysins over their non-iTSS counterparts. The slight improvement in PG degradation would represent an advantageous marginal gain of dual isoform Type I iTSS endolysins over their non-iTSS counterparts that may explain the prevalence of Type I iTSS endolysins across successive phage generations. Investigating this phenomenon can be challenging, particularly when studying it through the external application of recombinant endolysins; the kinetics of cleavage of particular bonds may be different, when the peptidoglycan is attacked from the cytoplasmic side compared to the extracellular side. Nonetheless, it is evident that Type I iTSS endolysins possess a distinct *in vivo* fitness advantage for the phage that, while subtle to discern under controlled laboratory conditions, is likely to manifest more prominently in natural environments.

Another interesting observation is that for the three-domain architecture typical for staphylococcal endolysins, iTSSs have only been identified for M23-Ami-CBD constructs and not for CHAP-Ami-CBD endolysins (Pinto et al. 2022). In contrast to Ply2638A, the amidase from CHAP-Ami-CBD endolysins seems to have only minimal bacteriolytic activity and has instead been suggested to play an auxiliary role via improving CBD binding to the target PG (Son et al. 2018). In the case of staphylococcal CHAP-Ami-CBD endolysins, it has been demonstrated that their bacteriolytic activity predominantly originates from the CHAP domain. This is evident in studies involving various staphylococcal endolysins, such as LysSA12 and LysSA97 (Son et al. 2018), LysGH15 (Gu et al. 2014), as well as LysK and φ11 (Navarre et al. 1999, Becker et al. 2009). Truncation studies, in which either the CHAP or amidase domain was independently fused to the native CBD, consistently revealed significantly greater activity associated with the CHAP domain compared to the amidase, which typically displayed reduced or even negligible activity in these investigations. Since, most enzymatic activity in CHAP-Ami-CBD endolysins seems to be associated with the CHAP domain, retaining an iTSS leading to co-expression of a low activity endolysin would not be beneficial and a waste of resources during phage infection. There are, however, other Gram-positive bacteria-targeting endolysins that feature a CHAP domain but no amidase domain and have an iTSS (Pinto et al. 2022). It is also important to note that CHAP-Ami-CBD endolysins are typically found by phages targeting *S. aureus*. However, as Ply2638A originates from a *S. pseudointermedius* phage, the observed differences in the use of iTSSs and amidase activity could simply be due to species variation.

In conclusion, the complexities of phage lysis and the development of multi-domain structures like Ply2638A endolysins offer a wealth of knowledge to explore. Here, we provide insights into the structural and functional attributes of Type I iTSS endolysin architectures that may prove instrumental in advancing the engineering of endolysins as precision antimicrobials in the future.

## Materials & Methods

### Bacterial strains and growth conditions

Bacterial strains used in this study are listed in Table 2. *E. coli* strains were grown at 37 °C in LB (10 g/L tryptone, 5 g/L yeast extract, 5 g/L NaCl, pH 7.8) or LB-PE (15 g/L tryptone, 8 g/L yeast extract, 6 g/L NaCl, pH 7.8) medium. All staphylococcal strains were grown in Brain Heart Infusion (BHI, Biolife Italiana) at 37 °C. *S. xylosus* L-forms used for phage engineering were grown in DM3 medium (5 g/L tryptone, 5 g/L yeast extract, 0.01 % BSA, 500 mM succinic acid, 5 g/L sucrose, 20 mM K_2_HPO_4_, 11 mM KH_2_PO_4_, 20mM MgCl_2_, pH 7.3) at 32 °C as described (Kilcher et al. 2018, Fernbach et al. 2024).

**Table 2.** Bacterial strains used in this study.

| STRAIN | RESISTANCE | APPLICATION | ORIGIN |
| --- | --- | --- | --- |
| *E. coli* BL21 Gold (DE3) | Tet<sup>R</sup> | Plasmid preparation, cloning, protein expression | Stratagene |
| *E. coli* XL-1BlueMRF' | Tet<sup>R</sup> | Plasmid preparation, cloning, protein expression | Stratagene |
| *S. aureus* Cowan I <sup>1)</sup> | MSSA | Activity assays | Clinical isolate (ATCC 12598) Bohacek et al. (1971) |
| *S. pseudointermedius* 2854<sup>2)</sup> | - | Phage propagation, phage assays | Slopek and Krzywy (1985) |
| *S. xylosus* L-form |  | Phage rebooting | Fernbach et al. (2024) |
| *S. aureus* BB270 (NCTC 8325 *mec*) |  | SH3b binding assays | Maidhof et al. (1991) |
| *S. aureus* BB270 (NCTC 8325 *mec* $\Delta femA$) | | SH3b binding assays | Maidhof et al. (1991) |
| *S. aureus* BB270 (NCTC 8325 *mec* $\Delta femB$) | | SH3b binding assays | Maidhof et al. (1991) |
| *S. aureus* BB270 (NCTC 8325 *mec* $\Delta femAB$) | | SH3b binding assays | Hubscher et al. (2007) |

<sup>1)</sup> Kindly provided by A. S. Zinkernagel (Bohacek et al. 1971)
<sup>2)</sup> Félix d'Hérelle Reference Center for Bacterial Viruses (Quebec City, QC, Canada)

### Phage engineering and production

Oligonucleotide pairs used for phage engineering are provided in SUPPLEMENTARY TABLE 1. Synthetic genomes were *in vitro* assembled using PCR-generated fragments and Gibson assembly (NEBuilder HiFi DNA Assembly Master Mix, BioLabs) with template DNA consisting of φ2638A WT gDNA that had been circularized through annealing of the terminal *cos* sites by heating gDNA to 65°C for 10 mins, followed by slow cooling at room temperature and ligation with T4 ligase (ThermoFisher). Circularized, synthetic genomes were dialyzed in distilled water and rebooted in *S. xylosus* Sul27 L-form cells as described (Kilcher et al. 2018, Fernbach et al. 2024). In brief, SuL27 L-forms were grown in DM3 medium supplemented with Penicillin G (200 µg/mL) and phosphomycin (500 µg/mL). After 48 h, the OD_600_ of the L-form culture was adjusted to 0.15 and the cells were mixed with the Gibson-assembled DNA or unmodified φ2638A gDNA (positive control), supplemented with 23 % (v/v) PEG 20,000 and incubated for 5 min at room temperature. The L-form transfection reaction was mixed with pre-warmed DM3 medium and assayed for mature phages after 24 h incubation at 37 °C by soft agar overlay using ½ BHI agar plates (37 g/L BHI, 12 g/L agar) and 5 mL BHI soft agar (37 g/L BHI, 6 g/L agar) spiked with 200 µL overnight grown *S. pseudintermedius* 2854 cells. Single plaques were picked and confirmed by PCR and Sanger sequencing (Microsynth, Switzerland).

### Phage propagation and purification

WT and engineered phages were propagated using soft agar overlays. 5 mL of soft BHI agar was spiked with 200 µL log-phase *S. pseudintermedius* 2854 cells and 10 µL of phages at ∼10^10^ PFU/mL and poured onto ½ BHI plates to produce semi-confluent lysis after overnight, 37°C incubation. Phage particles were washed out of the soft agar using 5Dml SM buffer per plate (100 mM NaCl, 8DmM MgSO_4_, and 50 mM Tris, pH 7.4) and filter-sterilized (0.2 μm) to obtain crude lysates. Lysates were further purified and concentrated by PEG precipitation (7 % PEG 8000 and 1DM NaCl), followed by CsCl isopycnic centrifugation and dialyzed twice against 1000× excess of SM buffer. The purified and concentrated phage stocks (∼10^12^ PFU/mL) were stored at 4 °C.

### Immunodetection of endolysin expression during phage infection

250 mL of exponentially growing cultures of *S. pseudintermedius* 2854 (OD_600_ ≈ 0.6, corresponding to 4.4 x 10^7^ CFU/mL) were infected with 250 µL of the phage stock leading to a final concentration of 4.4 x 10^7^ PFU/mL (MOI of 1) of φ2638A *ply*_WT-HA_, φ2638A *ply*_FL-HA_ and φ2638A *ply*_SV-HA_. Two samples of 1 mL were drawn from the culture at 30-minute intervals over 2 hours. One sample was used to measure the OD_600_, the other sample was centrifuged, OD-adjusted (OD_600_ = 10), and frozen (−20 °C) until analyzed. For SDS-PAGE analysis, 10 µL of the thawed bacterial suspensions were mixed with XT Sample Buffer (BioRad) supplemented with 50 mM DTT, heat denaturated at 100 °C for 10 min, and ran on TGX stain-free precast gels (Bio-Rad) for 40 min at 200 V. Proteins were transferred onto a PVDF membrane using an iBlot Gel Transfer System (Invitrogen). Western blotting was performed using an anti-HA mouse monoclonal antibody (Alexa Fluor® 488 anti-HA.11 Epitope Tag Antibody, BioLegend) as primary antibody diluted 1:1000 in TBS-T (20 mM Tris, 150 mM NaCl, 0.1 % Tween 20, pH 7.4) supplemented with 5 % BSA and HRP-conjugated rabbit anti-mouse antibody (Cell Signaling Technologies, USA) diluted 1:2000 as a secondary antibody.

### Recombinant protein construction, expression, and purification

Oligonucleotides, templates, and constructed plasmids are listed in SUPPLEMENTARY TABLE 2. Gene fragments were generated by PCR using φ2638A gDNA as template prior to NdeI, XhoI, or BamHI (NEB) restriction enzyme-based cloning into plasmids pET302 or pET200 depending on the construct. Construct Ply_FL_ was generated by site-directed mutagenesis of pET302_Ply_WT_ by mutating TTG>CTC at position Leu180 as previously described (Abaev et al. 2013). Individual plasmids were transformed into *E. coli* strains (Table 2) and grown in LB media supplemented with ampicillin (100 µg/mL; pET302 and pQE30) or kanamycin (50 µg/mL; pET200) at 37 °C until early log-phase. Cultures were cooled to 20 °C, induced with 0.5 mM isopropyl-β-D-thiogalactopyranoside (IPTG), and incubated for 18 h with agitation at 19 °C. Cells were harvested by centrifugation at 7,000 x g for 15 min, resuspended in Buffer A (20 mM Na_2_HPO_4_, 10 % glycerol, pH 7.4) for proteins without a His-tag and Lysis Buffer (50 mM Na_2_HPO_4_, 300 mM NaCl, 10 mM imidazole, 30 % glycerol, pH 8) for His-tagged proteins at 4 °C, and lysed using a Fluid Power Pressure Cell Homogenizer (Stansted). Proteins M23-2638A, GFP, GFP_SH3b2638A and GFP_SH3bLST harboring an N-terminal His-tag were purified by nickel affinity chromatography, as described elsewhere (Schmelcher et al. 2015). Ply_WT_, Ply_FL_, Ply_SV_, Ami and CBD were purified by cation exchange chromatography (CIEX) as follows: Cell extracts were centrifuged to remove cell debris at 20,000 x g for 60 min prior to loading of a 5 mL HiTrap SP-FF column on an ÄKTA purifier FPLC (GE Healthcare) equilibrated with Buffer A. Loaded extracts were washed with the running buffer for 5 column volumes (CVs) and eluted with Buffer B (20 mM Na_2_HPO_4_, 1 M NaCl, 10 % glycerol, pH 7.4) by applying a linear gradient (50 % in 10 CVs). All proteins underwent an additional purification step by size exclusion chromatography (SEC) on a HiLoad 16/60 Superdex 200 prep grade column (GE Healthcare) in SEC Buffer (50 mM Na_2_HPO_4_, 500 mM NaCl, 5 % glycerol, pH 7.4). Protein identity and purity were confirmed by SDS-PAGE, followed by Coomassie staining (InstantBlue^TM^, Sigma). Proteins were dialyzed into the following conditions: (i) For activity assays and SPR, PBS (10 mM Na_2_HPO_4_, 1.8 mM KH_2_PO_4_, 137 mM NaCl, 2.7 mM KCl, pH 7.4); (ii) for SEC-MALS, HEPES buffer (10 mM HEPES, 150 mM NaCl, 3.4 mM EDTA, 0.005 % Tween20, pH 7.4); (iii) for crystallization, TRIS buffer (20 mM Tris-HCl, 150 mM NaCl, pH 7.4).

### Turbidity reduction assays

For phage activity, an overnight culture of *S. pseudointermedius* 2854 was diluted 1:100 in fresh BHI, grown at 37 °C to OD_600nm_ ∼0.5, and then diluted in fresh BHI to OD_600nm_ = 0.1 (corresponding to ∼4 x 10^7^ CFU/mL). 100 µL of cells were added to a clear, flat bottom 96-well plate. Phage stocks were diluted in BHI and added to the cells at different titers (∼4 x 10^5^ to 10^9^ to PFU/mL) to infect at a range of MOIs from 0.001 to 100. OD_600nm_ was measured every 5 min over 18 h at 37 °C using a SPECTROstar Omega spectrophotometer (BMG Labtech) with shaking before every measurement. For turbidity reduction assays performed with endolysin complementation, phages were supplemented with purified, recombinant endolysin diluted in BHI prior to adding to the cells providing final concentrations of 10 nM, 100 nM, or 1 µM.

Recombinant endolysin activity was determined as described previously using premade and cryostocked *S. aureus* Cowan substrate cells (Abaev et al. 2013). Substrate cells were diluted to OD_600nm_ of 2 in PBS and incubated with serial dilutions of endolysins alone and in indicated molar ratios. Optical density was measured every 30 s over one hour. Lysis curves were normalized and control-corrected (cells alone). Data analysis was performed as previously described using a python script for automatization (Korndorfer et al. 2006, Abaev et al. 2013).

### Time-kill assays

An overnight culture of *S. pseudointermedius* 2854 was diluted 1:100 in fresh BHI, grown at 37 °C to OD_600nm_ ∼0.5, and then diluted in fresh BHI to OD_600nm_ = 0.1 (corresponding to ∼4 x 10^7^ CFU/mL). 500 µL of cells were added to a 1.5 mL Eppendorf tube. Phage stocks were diluted in BHI medium to titers between 4 x 10^7^ to 4 x 10^9^ PFU/mL with 500 µL added and mixed with the cells to reach MOIs between 1 to 100. At t=0, 50 µL of the negative control (i.e., cells mixed with phage-free BHI) was transferred to a dilution plate and 20 µL were serially diluted in PBS from 0 to 10^-7^. 20 µL were spotted on BHI agar square plates with the plate tilting method. Reaction tubes were incubated at 37 °C with shaking with 50 µL samples transferred to the dilution plate and 20 µL serially diluted and plated every hour for 6 hours as described above. CFUs were enumerated after overnight incubation at 37 °C.

### SEC coupled to Multiangle Light Scattering (SEC-MALS)

For SEC-MALS analysis 50 µL of protein samples of recombinant Ply_FL_, Ply_SV_ with varying ratios 1:1, 1:3 and 3:1 (w/w) at a concentration of 1 mg/mL were separated on a Superdex 200 10/30 column (GE Healthcare) using a LC1100 HPLC System (Agilent Technologies) coupled to an Optilab rEX refractometer (Wyatt Technology) and a miniDAWN three-angle light-scattering detector (Wyatt Technology). Protein separation was run in HEPES buffer (10 mM HEPES, 150 mM NaCl, 3.4 mM EDTA, 0.005 % Tween20, pH 7.4). Data analysis was performed using the ASTRA 6 software (Wyatt Technology).

### Surface Plasmon Resonance (SPR)

SPR was performed using a Biacore X system (GE Healthcare). The surface of a CMD500L chip (Xantec) was activated with 70 μL of a 1:1 ratio of 0.4 M 1-ethyl-3-(3-dimethylaminopropyl)-carbodiimide (EDC) and 0.1 M N-hydroxysuccinimide (NHS). 35 μL of Ply_FL_ (0.2 mg/mL) in immobilization buffer (10 mM sodium acetate, pH 4.2) was immobilized on the chip surface by amino coupling in flow cell 2 at a flow rate of 5 μL/min, according to the recommendations of the manufacturer (Biacore, GE Healthcare Life Sciences). Flow cell 1 was treated in the same way with EDC and NHS with no protein immobilized on the surface. Deactivation of the surface of both flow cells was done with 70 μL ethanolamine. 35 μL of the analytes (Ply_FL_, Ply_SV_, M23, Ami and CBD) at concentrations 750 nM, 1 μM, 2.5 μM and 5 μM (Supplementary Figure S7) were injected at a flow rate of 10 μL/min in running buffer (PBS: 10 mM Na_2_HPO_4_, 1.8 mM KH_2_PO_4_, 137 mM NaCl, 2.7 mM KCl, pH 7.4). Association and dissociation lasted 210 s and 720 s, respectively. Regeneration was performed by 5 μL injections of 10 mM glycine-HCl pH 2 between the samples. Baseline and injection point alignment were performed and the control-corrected sensorgrams were analyzed using the BiaEvaluation software (GE Healthcare).

### Fluorescence cell-binding assays

Binding of GFP-tagged CBDs to different *S. aureus* strains was assessed as previously described (Loessner et al. 2002). *S. aureus* BB270 wildtype and knockout strains Δ*femA*, Δ*femB* and Δ*femAB* were grown to early log phase. Cells were harvested by centrifugation and re-suspended in wash buffer (50 mM Na_2_HPO_4_, 120 mM NaCl, 0.1 % Tween20, pH 7.4) to an OD_600_ of 1.0. 20 µg of His- and GFP-tagged SH3b-2638A and SH3b-LST or His-tagged GFP (control) were added to 100 µL cells for 5 minutes, followed by washing twice with 0.5 mL wash buffer. Fluorescently labeled cells were resuspended in 50 µL and evaluated microscopically (Leica TCS SPE^TM^ CLSM) using a HCX PL FLUOTAR 100× oil objective and 15 % laser power (excitation 488 nm, emission 501-561 nm). The images were processed with Leica LAS AF Lite 3 software. In addition, fluorescence intensities of labeled cells were quantified (excitation 485 nm, emission 520 nm) using a FLUOstar Omega spectrophotometer (BMG LABTECH). Data sets were corrected against the GFP control, with binding values normalized to the *S. aureus* BB270 wildtype, which was set to 100 %.

### Protein crystallization

Crystallization screens were performed in 96-well format using the sitting-drop vapor-diffusion method at 20 °C using commercially available screens (Hampton Research, CA, USA; Molecular Dimensions, Suffolk, UK) with recombinant proteins in Tris buffer (20 mM Tris-HCl, 150 mM NaCl, pH 7.4) concentrated to 7-10 mg/mL. Crystals of the SH3b domain appeared with a reservoir composition of 1% (w/v) Tryptone, 0.05 M HEPES pH 7.0, 12 % (w/v) PEG-3350. Crystals of the M23 peptidase domain appeared with 0.2 M sodium chloride, 30 % (v/v) PEG-300, pH 5.7. Larger crystals were produced after optimization of the conditions for subsequent hanging-drop vapor-diffusion crystallization. The crystals were grown at 19°C in hanging drops containing 1 μl protein solution (10 mg/mL for SH3b and 7 mg/mL for M23 peptidase) and 1 μl crystallization solution (1 % (w/v) Tryptone, 0.05 M HEPES pH 7.0, 14 % (w/v) PEG 3350 for SH3b and 0.2 M sodium chloride, 28 % (v/v) PEG 300, pH 5.7 for M23 peptidase), against a 1 ml reservoir crystallization solution. Crystals were fished and cryo-protected in the same crystallization solutions containing 30 % glycerol.

### X-ray crystallography data collection and refinement

X-ray diffraction data was collected on the X06SA (PXI) beamline at the Swiss Light Source, Paul Scherrer Institute, Switzerland, using an Eiger-16M X (DECTRIS Ltd., Baden-Dättwil, Germany) pixel detector at 100 K and wavelength 1.00 Å. Single datasets were collected and indexed, integrated, and scaled using XDS (Kabsch 2010). For the M23 peptidase domain, a single dataset was collected to 2.3 Å resolution in the space group P2_1_. The structure was solved by molecular replacement using Phaser (McCoy et al. 2007) and a hybrid search model built from the enzymatic domains of various peptidase domains using Phenix.sculptor with two molecules identified in the asymmetric unit. For the CBD, a single dataset was collected to 2.8 Å resolution, indexed, integrated and scaled using XDS with space group P1 (Kabsch 2010). Cell content analysis suggested 7 to 10 molecules per asymmetric unit with eight molecules providing a Matthew’s coefficient of 2.71 Å^3^/Da and 54.7 % solvent content. The structure was solved by molecular replacement using MolRep (Vagin and Teplyakov 1997) and the CBD of lysostaphin (PDB ID: 5LEO) as a search model with a final eight molecules identified within the asymmetric unit. For all constructs, successive rounds of refinement were performed using phenix.refine (Afonine et al. 2012) and Coot (Emsley et al. 2010) to generate final models that were validated using MolProbity (Chen et al. 2010). The CheckMyMetal webserver (Zheng et al. 2017) was used to validate the coordination geometry of all metal ions. The DALI server was used to identify structural homologs in the PDB (Holm et al. 2023).

All structure figures were created using PyMOL version 2.5.3 (Schrodinger LLC) with electrostatic surface potential calculated using the Adaptive Poisson-Boltzmann Solver APBS plugin (Baker et al. 2001). Crystallographic data collection and refinement statistics are provided in Table 1. Structures and X-ray diffraction data were deposited at the PDB under the accession codes 6YJ1 and 7AQH for the Ply2638A M23 peptidase domain and SH3b CBD, respectively.

### Structure prediction and analysis

Structure predictions were performed using AlphaFold 2.0 (Jumper et al. 2021) and AlphaFold-Multimer (Evans et al. 2022) downloaded from www.github.com/deepmind/alphafold and installed on a HP Z6 workstation equipped with a Xeon Gold 6354 CPU, 192 GB of RAM, an Nvidia RTX 2080TI GPU, and M2 SSD disks, running Ubuntu Linux 20.04. All predictions were assessed using internally generated confidence scores. Confidence per residue is provided as a predicted Local Distance Difference Test score (pLDDT; scored 0-100), with the average of all residues per model provided in the main text and figure legends. A pLDDT ≥ 90 have very high model confidence, residues with 90 > pLDDT ≥ 70 are classified as confident, while residues with 70 > pLDDT > 50 have low confidence.

### Statistical analyses

All statistical analyses were conducted using GraphPad Prism 8.2.0 (GraphPad Software, San Diego, CA, United States). For the phage infection assays, comparisons between engineered and wildtype φ2638A phages were performed using unpaired Student’s t-tests. One-way ANOVA, followed by Dunnett’s multiple comparisons test, was employed to assess the differences among Ply_FL_, Ply_SV_, and their combinations in TRAs. The relative binding of GFP-tagged SH3b domains to S. aureus cells was assessed using unpaired Student’s t-tests. A significance level of p < 0.05 was considered statistically significant.

## Acknowledgements

This work was supported by funds from Micreos BV, The Hague, The Netherlands (to M.J.L.). The funders had no role in study design, data collection and interpretation, or the decision to submit the work for publication. The authors would like to acknowledge Eric Sumrall and Yang Shen (ETH Zurich) for their assistance in preparation of the Western Blots and SPR data, respectively. We would also like to thank Beat Blattmann from the Protein Crystallization Center (PCC) at University of Zurich, and the staff from the X06SA beamline of the Swiss Light Source (Paul Scherrer Institute, Villigen). We would like to thank Raphael Haemmerli and Elia Haemmerli for their contribution to the python code for analysis of the protein TRAs.

## Competing interests

S.K., A.K., M.S., and M.D. are employees of Micreos Pharmaceuticals, a company developing bacteriophage- and endolysin-based antimicrobials. M.J.L. serves as a scientific advisor to Micreos Pharmaceuticals. The remaining authors declare no conflicts of interest. All research presented in this paper was conducted at ETH Zurich and the University of Zurich.

## Contributions

Conceptualization, L.V.Z., A.M.S., M.S., M.D., Methodology, L.V.Z., A.M.S., M.D.; Project administration, M.D; Investigation, L.V.Z., A.M.S., P.E., S.M., S.K., C.I., A.K., B.D., P.R.E.M., M.D.; X-ray data collection and analysis, P.E., P.R.E.M., M.D.; Data curation, L.V.Z., A.M.S., M.D.; Visualization, L.V.Z., A.M.S., M.D.; Writing—manuscript, L.V.Z., M.D.; Writing—review and editing, L.V.Z., A.M.S., P.E., B.D., P.R.E.M, A.P., M.J.L, M.S., M.D.; Funding acquisition, M.J.L., M.S., M.D.; Resources, M.J.L., P.R.E.M., A.P., M.J.L., M.D.

## Notes

https://www.rcsb.org/structure/6yj1

https://www.rcsb.org/structure/7AQH

## References

Abaev, I., J. Foster-Frey, O. Korobova, N. Shishkova, N. Kiseleva, P. Kopylov, S. Pryamchuk, M. Schmelcher, S. C. Becker and D. M. Donovan (2013). “Staphylococcal phage 2638A endolysin is lytic for *Staphylococcus aureus* and harbors an inter-lytic-domain secondary translational start site.” Appl Microbiol Biotechnol 97(8): 3449–3456.

Afonine, P. V., R. W. Grosse-Kunstleve, N. Echols, J. J. Headd, N. W. Moriarty, M. Mustyakimov, T. C. Terwilliger, A. Urzhumtsev, P. H. Zwart and P. D. Adams (2012). “Towards automated crystallographic structure refinement with phenix.refine.” Acta Crystallogr D Biol Crystallogr 68(Pt 4): 352–367.

An, D. R., H. S. Kim, J. Kim, H. N. Im, H. J. Yoon, J. Y. Yoon, J. Y. Jang, D. Hesek, M. Lee, S. Mobashery, S. J. Kim, B. I. Lee and S. W. Suh (2015). “Structure of Csd3 from Helicobacter pylori, a cell shape-determining metallopeptidase.” Acta Crystallogr D Biol Crystallogr 71(Pt 3): 675–686.

Ashkenazy, H., S. Abadi, E. Martz, O. Chay, I. Mayrose, T. Pupko and N. Ben-Tal (2016). “ConSurf 2016: an improved methodology to estimate and visualize evolutionary conservation in macromolecules.” Nucleic Acids Res 44(W1): W344–350.

Baker, N. A., D. Sept, S. Joseph, M. J. Holst and J. A. McCammon (2001). “Electrostatics of nanosystems: Application to microtubules and the ribosome.” Proceedings of the National Academy of Sciences 98(18): 10037–10041.

Becker, S. C., S. Dong, J. R. Baker, J. Foster-Frey, D. G. Pritchard and D. M. Donovan (2009). “LysK CHAP endopeptidase domain is required for lysis of live staphylococcal cells.” FEMS Microbiology Letters 294(1): 52–60.

Becker, S. C., J. Foster-Frey, A. J. Stodola, D. Anacker and D. M. Donovan (2009). “Differentially conserved staphylococcal SH3b_5 cell wall binding domains confer increased staphylolytic and streptolytic activity to a streptococcal prophage endolysin domain.” Gene 443(1-2): 32–41.

Bohacek, J., M. Kocur and T. Martinec (1971). “Deoxyribonucleic acid base composition of serotype strains of Staphylococcus aureus.” J Gen Microbiol 68(1): 109–113.

Büttner, F. M., S. Zoll, M. Nega, F. Götz and T. Stehle (2014). “Structure-Function Analysis of Staphylococcus aureus Amidase Reveals the Determinants of Peptidoglycan Recognition and Cleavage*.” Journal of Biological Chemistry 289(16): 11083–11094.

Callaway, E. (2022). “What’s next for AlphaFold and the AI protein-folding revolution.” Nature 604(7905): 234–238.

Catalao, M. J., C. Milho, F. Gil, J. Moniz-Pereira and M. Pimentel (2011). “A second endolysin gene is fully embedded in-frame with the lysA gene of mycobacteriophage Ms6.” PLoS One 6(6): e20515.

Chen, V. B., W. B. Arendall, 3rd, J. J. Headd, D. A. Keedy, R. M. Immormino, G. J. Kapral, L. W. Murray, J. S. Richardson and D. C. Richardson (2010). “MolProbity: all-atom structure validation for macromolecular crystallography.” Acta Crystallogr D Biol Crystallogr 66(Pt 1): 12–21.

Cheng, X., X. Zhang, J. W. Pflugrath and F. W. Studier (1994). “The structure of bacteriophage T7 lysozyme, a zinc amidase and an inhibitor of T7 RNA polymerase.” Proceedings of the National Academy of Sciences 91(9): 4034–4038.

Danis-Wlodarczyk, K. M., D. J. Wozniak and S. T. Abedon (2021). “Treating Bacterial Infections with Bacteriophage-Based Enzybiotics: In Vitro, In Vivo and Clinical Application.” Antibiotics 10(12): 1497.

Diaz, E., R. Lopez and J. L. Garcia (1990). “Chimeric phage-bacterial enzymes: a clue to the modular evolution of genes.” Proc Natl Acad Sci U S A 87(20): 8125–8129.

Dunne, M., S. Leicht, B. Krichel, H. D. Mertens, A. Thompson, J. Krijgsveld, D. I. Svergun, N. Gomez-Torres, S. Garde, C. Uetrecht, A. Narbad, M. J. Mayer and R. Meijers (2016). “Crystal Structure of the CTP1L Endolysin Reveals How Its Activity Is Regulated by a Secondary Translation Product.” J Biol Chem 291(10): 4882–4893.

Eichenseher, F., B. L. Herpers, P. Badoux, J. M. Leyva-Castillo, R. S. Geha, M. van der Zwart, J. McKellar, F. Janssen, B. de Rooij, L. Selvakumar, C. Rohrig, J. Frieling, M. Offerhaus, M. J. Loessner and M. Schmelcher (2022). “Linker-Improved Chimeric Endolysin Selectively Kills Staphylococcus aureus In Vitro, on Reconstituted Human Epidermis, and in a Murine Model of Skin Infection.” Antimicrob Agents Chemother 66(5): e0227321.

Emsley, P., B. Lohkamp, W. G. Scott and K. Cowtan (2010). “Features and development of Coot.” Acta Crystallographica Section D: Biological Crystallography 66(4): 486–501.

Evans, R., M. O’Neill, A. Pritzel, N. Antropova, A. Senior, T. Green, A. Žídek, R. Bates, S. Blackwell, J. Yim, O. Ronneberger, S. Bodenstein, M. Zielinski, A. Bridgland, A. Potapenko, A. Cowie, K. Tunyasuvunakool, R. Jain, E. Clancy, P. Kohli, J. Jumper and D. Hassabis (2022). “Protein complex prediction with AlphaFold-Multimer.” bioRxiv: 2021.2010.2004.463034.

Fernbach, J., S. Meile, S. Kilcher and M. J. Loessner (2024). “Genetic Engineering and Rebooting of Bacteriophages in L-Form Bacteria.” Methods Mol Biol 2734: 247–259.

Fischetti, V. A. (2010). “Bacteriophage endolysins: a novel anti-infective to control Gram-positive pathogens.” International Journal of Medical Microbiology 300(6): 357–362.

Fowler, V. G., A. F. Das, J. Lipka-Diamond, R. Schuch, R. Pomerantz, L. Jáuregui-Peredo, A. Bressler, D. Evans, G. J. Moran and M. E. Rupp (2020). “Exebacase for patients with *Staphylococcus aureus* bloodstream infection and endocarditis.” The Journal of clinical investigation 130(7): 3750–3760.

Gonzalez-Delgado, L. S., H. Walters-Morgan, B. Salamaga, A. J. Robertson, A. M. Hounslow, E. Jagielska, I. Sabała, M. P. Williamson, A. L. Lovering and S. Mesnage (2020). “Two-site recognition of Staphylococcus aureus peptidoglycan by lysostaphin SH3b.” Nature chemical biology 16(1): 24–30.

Gonzalez-Serrano, R., R. Rosselli, J. J. Roda-Garcia, A.-B. Martin-Cuadrado, F. Rodriguez-Valera and M. Dunne (2023). “Distantly related Alteromonas bacteriophages share tail fibers exhibiting properties of transient chaperone caps.” Nature Communications 14(1): 6517.

Grabowska, M., E. Jagielska, H. Czapinska, M. Bochtler and I. Sabala (2015). “High resolution structure of an M23 peptidase with a substrate analogue.” Scientific Reports 5(1): 14833.

Gu, J., Y. Feng, X. Feng, C. Sun, L. Lei, W. Ding, F. Niu, L. Jiao, M. Yang, Y. Li, X. Liu, J. Song, Z. Cui, D. Han, C. Du, Y. Yang, S. Ouyang, Z. J. Liu and W. Han (2014). “Structural and biochemical characterization reveals LysGH15 as an unprecedented “EF-hand-like” calcium-binding phage lysin.” PLoS Pathog 10(5): e1004109.

Gu, J., W. Xu, L. Lei, J. Huang, X. Feng, C. Sun, C. Du, J. Zuo, Y. Li, T. Du, L. Li and W. Han (2011). “LysGH15, a novel bacteriophage lysin, protects a murine bacteremia model efficiently against lethal methicillin-resistant *Staphylococcus aureus* infection.” J Clin Microbiol 49(1): 111–117.

Gutierrez, D., P. Ruas-Madiedo, B. Martinez, A. Rodriguez and P. Garcia (2014). “Effective removal of staphylococcal biofilms by the endolysin LysH5.” PLoS One 9(9): e107307.

Haddad Kashani, H., M. Schmelcher, H. Sabzalipoor, E. Seyed Hosseini and R. Moniri (2018). “Recombinant Endolysins as Potential Therapeutics against Antibiotic-Resistant Staphylococcus aureus: Current Status of Research and Novel Delivery Strategies.” Clin Microbiol Rev 31(1).

Holm, L., A. Laiho, P. Törönen and M. Salgado (2023). “DALI shines a light on remote homologs: One hundred discoveries.” Protein Sci 32(1): e4519.

Hubscher, J., A. Jansen, O. Kotte, J. Schafer, P. A. Majcherczyk, L. G. Harris, G. Bierbaum, M. Heinemann and B. Berger-Bachi (2007). “Living with an imperfect cell wall: compensation of femAB inactivation in Staphylococcus aureus.” BMC Genomics 8: 307.

Humphreys, I. R., J. Pei, M. Baek, A. Krishnakumar, I. Anishchenko, S. Ovchinnikov, J. Zhang, T. J. Ness, S. Banjade, S. R. Bagde, V. G. Stancheva, X. H. Li, K. Liu, Z. Zheng, D. J. Barrero, U. Roy, J. Kuper, I. S. Fernández, B. Szakal, D. Branzei, J. Rizo, C. Kisker, E. C. Greene, S. Biggins, S. Keeney, E. A. Miller, J. C. Fromme, T. L. Hendrickson, Q. Cong and D. Baker (2021). “Computed structures of core eukaryotic protein complexes.” Science 374(6573): eabm4805.

Jumper, J., R. Evans, A. Pritzel, T. Green, M. Figurnov, O. Ronneberger, K. Tunyasuvunakool, R. Bates, A. Zidek, A. Potapenko, A. Bridgland, C. Meyer, S. A. A. Kohl, A. J. Ballard, A. Cowie, B. Romera-Paredes, S. Nikolov, R. Jain, J. Adler, T. Back, S. Petersen, D. Reiman, E. Clancy, M. Zielinski, M. Steinegger, M. Pacholska, T. Berghammer, S. Bodenstein, D. Silver, O. Vinyals, A. W. Senior, K. Kavukcuoglu, P. Kohli and D. Hassabis (2021). “Highly accurate protein structure prediction with AlphaFold.” Nature 596(7873): 583–589.

Jun, S. Y., G. M. Jung, S. J. Yoon, Y. J. Choi, W. S. Koh, K. S. Moon and S. H. Kang (2014). “Preclinical safety evaluation of intravenously administered SAL200 containing the recombinant phage endolysin SAL-1 as a pharmaceutical ingredient.” Antimicrob Agents Chemother 58(4): 2084–2088.

Kabsch, W. (2010). “Xds.” Acta Crystallogr D Biol Crystallogr 66(Pt 2): 125–132.

Kilcher, S., P. Studer, C. Muessner, J. Klumpp and M. J. Loessner (2018). “Cross-genus rebooting of custom-made, synthetic bacteriophage genomes in L-form bacteria.” Proc Natl Acad Sci U S A 115(3): 567–572.

Korndorfer, I. P., J. Danzer, M. Schmelcher, M. Zimmer, A. Skerra and M. J. Loessner (2006). “The crystal structure of the bacteriophage PSA endolysin reveals a unique fold responsible for specific recognition of Listeria cell walls.” J Mol Biol 364(4): 678–689.

Kuiper, J. W. P., J. M. A. Hogervorst, B. L. Herpers, A. D. Bakker, J. Klein-Nulend, P. A. Nolte and B. P. Krom (2021). “The novel endolysin XZ. 700 effectively treats MRSA biofilms in two biofilm models without showing toxicity on human bone cells in vitro.” Biofouling 37(2): 184–193.

Lee, C. Y., D. Hubrich, J. K. Varga, C. Schäfer, M. Welzel, E. Schumbera, M. Đokić, J. M. Strom, J. Schönfeld, J. L. Geist, F. Polat, T. J. Gibson, C. I. K. Valsecchi, M. Kumar, O. Schueler-Furman and K. Luck (2023). “Systematic discovery of protein interaction interfaces using AlphaFold and experimental validation.” bioRxiv: 2023.2008.2007.552219.

Loessner, M. J., S. Gaeng, G. Wendlinger, S. K. Maier and S. Scherer (1998). “The two-component lysis system of *Staphylococcus aureus* bacteriophage Twort: a large TTG-start holin and an associated amidase endolysin.” FEMS Microbiol Lett 162(2): 265–274.

Loessner, M. J., K. Kramer, F. Ebel and S. Scherer (2002). “C-terminal domains of Listeria monocytogenes bacteriophage murein hydrolases determine specific recognition and high-affinity binding to bacterial cell wall carbohydrates.” Mol Microbiol 44(2): 335–349.

Low, L. Y., C. Yang, M. Perego, A. Osterman and R. Liddington (2011). “Role of Net Charge on Catalytic Domain and Influence of Cell Wall Binding Domain on Bactericidal Activity, Specificity, and Host Range of Phage Lysins*.” Journal of Biological Chemistry 286(39): 34391–34403.

Low, L. Y., C. Yang, M. Perego, A. Osterman and R. C. Liddington (2005). “Structure and Lytic Activity of a Bacillus anthracis Prophage Endolysin*.” Journal of Biological Chemistry 280(42): 35433–35439.

Lu, Y., Y. Wang, J. Wang, Y. Zhao, Q. Zhong, G. Li, Z. Fu and S. Lu (2021). “Phage Endolysin LysP108 Showed Promising Antibacterial Potential Against Methicillin-resistant Staphylococcus aureus.” Front Cell Infect Microbiol 11: 668430.

Maidhof, H., B. Reinicke, P. Blumel, B. Berger-Bachi and H. Labischinski (1991). “femA, which encodes a factor essential for expression of methicillin resistance, affects glycine content of peptidoglycan in methicillin-resistant and methicillin-susceptible Staphylococcus aureus strains.” J Bacteriol 173(11): 3507–3513.

Małecki, P. H., P. Mitkowski, E. Jagielska, K. Trochimiak, S. Mesnage and I. Sabała (2021). “Structural Characterization of EnpA D,L-Endopeptidase from Enterococcus faecalis Prophage Provides Insights into Substrate Specificity of M23 Peptidases.” Int J Mol Sci 22(13).

McCoy, A. J., R. W. Grosse-Kunstleve, P. D. Adams, M. D. Winn, L. C. Storoni and R. J. Read (2007). “Phaser crystallographic software.” J Appl Crystallogr 40(Pt 4): 658–674.

Mitkowski, P., E. Jagielska, E. Nowak, J. M. Bujnicki, F. Stefaniak, D. Niedziałek, M. Bochtler and I. Sabała (2019). “Structural bases of peptidoglycan recognition by lysostaphin SH3b domain.” Scientific reports 9(1): 5965.

Murray, C. J. L., K. S. Ikuta, F. Sharara, L. Swetschinski, G. Robles Aguilar, A. Gray, C. Han, C. Bisignano, P. Rao, E. Wool, S. C. Johnson, A. J. Browne, M. G. Chipeta, F. Fell, S. Hackett, G. Haines-Woodhouse, B. H. Kashef Hamadani, E. A. P. Kumaran, B. McManigal, S. Achalapong, R. Agarwal, S. Akech, S. Albertson, J. Amuasi, J. Andrews, A. Aravkin, E. Ashley, F.-X. Babin, F. Bailey, S. Baker, B. Basnyat, A. Bekker, R. Bender, J. A. Berkley, A. Bethou, J. Bielicki, S. Boonkasidecha, J. Bukosia, C. Carvalheiro, C. Castañeda-Orjuela, V. Chansamouth, S. Chaurasia, S. Chiurchiù, F. Chowdhury, R. Clotaire Donatien, A. J. Cook, B. Cooper, T. R. Cressey, E. Criollo-Mora, M. Cunningham, S. Darboe, N. P. J. Day, M. De Luca, K. Dokova, A. Dramowski, S. J. Dunachie, T. Duong Bich, T. Eckmanns, D. Eibach, A. Emami, N. Feasey, N. Fisher-Pearson, K. Forrest, C. Garcia, D. Garrett, P. Gastmeier, A. Z. Giref, R. C. Greer, V. Gupta, S. Haller, A. Haselbeck, S. I. Hay, M. Holm, S. Hopkins, Y. Hsia, K. C. Iregbu, J. Jacobs, D. Jarovsky, F. Javanmardi, A. W. J. Jenney, M. Khorana, S. Khusuwan, N. Kissoon, E. Kobeissi, T. Kostyanev, F. Krapp, R. Krumkamp, A. Kumar, H. H. Kyu, C. Lim, K. Lim, D. Limmathurotsakul, M. J. Loftus, M. Lunn, J. Ma, A. Manoharan, F. Marks, J. May, M. Mayxay, N. Mturi, T. Munera-Huertas, P. Musicha, L. A. Musila, M. M. Mussi-Pinhata, R. N. Naidu, T. Nakamura, R. Nanavati, S. Nangia, P. Newton, C. Ngoun, A. Novotney, D. Nwakanma, C. W. Obiero, T. J. Ochoa, A. Olivas-Martinez, P. Olliaro, E. Ooko, E. Ortiz-Brizuela, P. Ounchanum, G. D. Pak, J. L. Paredes, A. Y. Peleg, C. Perrone, T. Phe, K. Phommasone, N. Plakkal, A. Ponce-de-Leon, M. Raad, T. Ramdin, S. Rattanavong, A. Riddell, T. Roberts, J. V. Robotham, A. Roca, V. D. Rosenthal, K. E. Rudd, N. Russell, H. S. Sader, W. Saengchan, J. Schnall, J. A. G. Scott, S. Seekaew, M. Sharland, M. Shivamallappa, J. Sifuentes-Osornio, A. J. Simpson, N. Steenkeste, A. J. Stewardson, T. Stoeva, N. Tasak, A. Thaiprakong, G. Thwaites, C. Tigoi, C. Turner, P. Turner, H. R. van Doorn, S. Velaphi, A. Vongpradith, M. Vongsouvath, H. Vu, T. Walsh, J. L. Walson, S. Waner, T. Wangrangsimakul, P. Wannapinij, T. Wozniak, T. E. M. W. Young Sharma, K. C. Yu, P. Zheng, B. Sartorius, A. D. Lopez, A. Stergachis, C. Moore, C. Dolecek and M. Naghavi (2022). “Global burden of bacterial antimicrobial resistance in 2019: a systematic analysis.” The Lancet 399(10325): 629–655.

Navarre, W. W., H. Ton-That, K. F. Faull and O. Schneewind (1999). “Multiple enzymatic activities of the murein hydrolase from staphylococcal phage phi11. Identification of a D-alanyl-glycine endopeptidase activity.” J Biol Chem 274(22): 15847–15856.

O’Flaherty, S., A. Coffey, W. Meaney, G. F. Fitzgerald and R. P. Ross (2005). “The recombinant phage lysin LysK has a broad spectrum of lytic activity against clinically relevant staphylococci, including methicillin-resistant Staphylococcus aureus.” J Bacteriol 187(20): 7161–7164.

Olsen, N. M. C., E. Thiran, T. Hasler, T. Vanzieleghem, G. N. Belibasakis, J. Mahillon, M. J. Loessner and M. Schmelcher (2018). “Synergistic Removal of Static and Dynamic Staphylococcus aureus Biofilms by Combined Treatment with a Bacteriophage Endolysin and a Polysaccharide Depolymerase.” Viruses 10(8).

Pallesen, E. M. H., M. Gluud, C. K. Vadivel, T. B. Buus, B. de Rooij, Z. Zeng, S. Ahmad, A. Willerslev-Olsen, C. Röhrig, M. R. Kamstrup, L. Bay, L. Lindahl, T. Krejsgaard, C. Geisler, C. M. Bonefeld, L. Iversen, A. Woetmann, S. B. Koralov, T. Bjarnsholt, J. Frieling, M. Schmelcher and N. Ødum (2023). “Endolysin Inhibits Skin Colonization by Patient-Derived Staphylococcus Aureus and Malignant T-Cell Activation in Cutaneous T-Cell Lymphoma.” J Invest Dermatol 143(9): 1757–1768.e1753.

Pavlopoulos, G. A., F. A. Baltoumas, S. Liu, O. Selvitopi, A. P. Camargo, S. Nayfach, A. Azad, S. Roux, L. Call, N. N. Ivanova, I. M. Chen, D. Paez-Espino, E. Karatzas, S. G. Acinas, N. Ahlgren, G. Attwood, P. Baldrian, T. Berry, J. M. Bhatnagar, D. Bhaya, K. D. Bidle, J. L. Blanchard, E. S. Boyd, J. L. Bowen, J. Bowman, S. H. Brawley, E. L. Brodie, A. Brune, D. A. Bryant, A. Buchan, H. Cadillo-Quiroz, B. J. Campbell, R. Cavicchioli, P. F. Chuckran, M. Coleman, S. Crowe, D. R. Colman, C. R. Currie, J. Dangl, N. Delherbe, V. J. Denef, P. Dijkstra, D. D. Distel, E. Eloe-Fadrosh, K. Fisher, C. Francis, A. Garoutte, A. Gaudin, L. Gerwick, F. Godoy-Vitorino, P. Guerra, J. Guo, M. Y. Habteselassie, S. J. Hallam, R. Hatzenpichler, U. Hentschel, M. Hess, A. M. Hirsch, L. A. Hug, J. Hultman, D. E. Hunt, M. Huntemann, W. P. Inskeep, T. Y. James, J. Jansson, E. R. Johnston, M. Kalyuzhnaya, C. N. Kelly, R. M. Kelly, J. L. Klassen, K. Nüsslein, J. E. Kostka, S. Lindow, E. Lilleskov, M. Lynes, R. Mackelprang, F. M. Martin, O. U. Mason, R. M. McKay, K. McMahon, D. A. Mead, M. Medina, L. K. Meredith, T. Mock, W. W. Mohn, M. A. Moran, A. Murray, J. D. Neufeld, R. Neumann, J. M. Norton, L. P. Partida-Martinez, N. Pietrasiak, D. Pelletier, T. B. K. Reddy, B. K. Reese, N. J. Reichart, R. Reiss, M. A. Saito, D. P. Schachtman, R. Seshadri, A. Shade, D. Sherman, R. Simister, H. Simon, J. Stegen, R. Stepanauskas, M. Sullivan, D. Y. Sumner, H. Teeling, K. Thamatrakoln, K. Treseder, S. Tringe, P. Vaishampayan, D. L. Valentine, N. B. Waldo, M. P. Waldrop, D. A. Walsh, D. M. Ward, M. Wilkins, T. Whitman, J. Woolet, T. Woyke, I. Iliopoulos, K. Konstantinidis, J. M. Tiedje, J. Pett-Ridge, D. Baker, A. Visel, C. A. Ouzounis, S. Ovchinnikov, A. Buluç, N. C. Kyrpides and C. Novel Metagenome Protein Families (2023). “Unraveling the functional dark matter through global metagenomics.” Nature 622(7983): 594–602.

Pingoud, A. and A. Jeltsch (2001). “Structure and function of type II restriction endonucleases.” Nucleic Acids Res 29(18): 3705–3727.

Pinto, D., R. Goncalo, M. Louro, M. S. Silva, G. Hernandez, T. N. Cordeiro, C. Cordeiro and C. Sao-Jose (2022). “On the Occurrence and Multimerization of Two-Polypeptide Phage Endolysins Encoded in Single Genes.” Microbiol Spectr 10(4): e0103722.

Proenca, D., C. Velours, C. Leandro, M. Garcia, M. Pimentel and C. Sao-Jose (2015). “A two-component, multimeric endolysin encoded by a single gene.” Mol Microbiol 95(5): 739–753.

Rahman, M. U., W. Wang, Q. Sun, J. A. Shah, C. Li, Y. Sun, Y. Li, B. Zhang, W. Chen and S. Wang (2021). “Endolysin, a Promising Solution against Antimicrobial Resistance.” Antibiotics (Basel) 10(11).

Razew, A., J. N. Schwarz, P. Mitkowski, I. Sabala and M. Kaus-Drobek (2022). “One fold, many functions-M23 family of peptidoglycan hydrolases.” Front Microbiol 13: 1036964.

Sabala, I., E. Jagielska, P. T. Bardelang, H. Czapinska, S. O. Dahms, J. A. Sharpe, R. James, M. E. Than, N. R. Thomas and M. Bochtler (2014). “Crystal structure of the antimicrobial peptidase lysostaphin from Staphylococcus simulans.” The FEBS journal 281(18): 4112–4122.

Sanz-Gaitero, M., R. Keary, C. Garcia-Doval, A. Coffey and M. J. van Raaij (2014). “Crystal structure of the lytic CHAP(K) domain of the endolysin LysK from Staphylococcus aureus bacteriophage K.” Virol J 11: 133.

Sayers, E. W., E. E. Bolton, J. R. Brister, K. Canese, J. Chan, D. C. Comeau, R. Connor, K. Funk, C. Kelly, S. Kim, T. Madej, A. Marchler-Bauer, C. Lanczycki, S. Lathrop, Z. Lu, F. Thibaud-Nissen, T. Murphy, L. Phan, Y. Skripchenko, T. Tse, J. Wang, R. Williams, B. W. Trawick, K. D. Pruitt and S. T. Sherry (2022). “Database resources of the national center for biotechnology information.” Nucleic Acids Res 50(D1): D20–d26.

Schmelcher, M., D. M. Donovan and M. J. Loessner (2012). “Bacteriophage endolysins as novel antimicrobials.” Future Microbiol 7(10): 1147–1171.

Schmelcher, M. and M. J. Loessner (2021). “Bacteriophage endolysins—extending their application to tissues and the bloodstream.” Current opinion in biotechnology 68: 51–59.

Schmelcher, M., Y. Shen, D. C. Nelson, M. R. Eugster, F. Eichenseher, D. C. Hanke, M. J. Loessner, S. Dong, D. G. Pritchard, J. C. Lee, S. C. Becker, J. Foster-Frey and D. M. Donovan (2015). “Evolutionarily distinct bacteriophage endolysins featuring conserved peptidoglycan cleavage sites protect mice from MRSA infection.” J Antimicrob Chemother 70(5): 1453–1465.

Schuch, R., H. M. Lee, B. C. Schneider, K. L. Sauve, C. Law, B. K. Khan, J. A. Rotolo, Y. Horiuchi, D. E. Couto, A. Raz, V. A. Fischetti, D. B. Huang, R. C. Nowinski and M. Wittekind (2014). “Combination therapy with lysin CF-301 and antibiotic is superior to antibiotic alone for treating methicillin-resistant Staphylococcus aureus-induced murine bacteremia.” J Infect Dis 209(9): 1469–1478.

Slopek, S. and T. Krzywy (1985). “Morphology and ultrastructure of bacteriophages. An electron microscopic study.” Arch Immunol Ther Exp (Warsz) 33(1): 1–217.

Sobral, R. and A. Tomasz (2019). “The Staphylococcal Cell Wall.” Microbiol Spectr 7(4).

Son, B., M. Kong, Y. Lee and S. Ryu (2020). “Development of a Novel Chimeric Endolysin, Lys109 With Enhanced Lytic Activity Against Staphylococcus aureus.” Front Microbiol 11: 615887.

Son, B., M. Kong and S. Ryu (2018). “The Auxiliary Role of the Amidase Domain in Cell Wall Binding and Exolytic Activity of Staphylococcal Phage Endolysins.” Viruses 10(6).

Vagin, A. and A. Teplyakov (1997). “MOLREP: an automated program for molecular replacement.” Journal of applied crystallography 30(6): 1022–1025.

Vollmer, W. and S. J. Seligman (2010). “Architecture of peptidoglycan: more data and more models.” Trends in Microbiology 18(2): 59–66.

WHO (2020). “Global antimicrobial resistance surveillance system (GLASS) report: early implementation 2020.”

Xu, X., D. Zhang, B. Zhou, X. Zhen and S. Ouyang (2021). “Structural and biochemical analyses of the tetrameric cell binding domain of Lys170 from enterococcal phage F170/08.” European Biophysics Journal 50(5): 721–729.

Zheng, H., D. R. Cooper, P. J. Porebski, I. G. Shabalin, K. B. Handing and W. Minor (2017). “CheckMyMetal: a macromolecular metal-binding validation tool.” Acta Crystallogr D Struct Biol 73(Pt 3): 223–233.

Zhou, B., X. Zhen, H. Zhou, F. Zhao, C. Fan, V. Perculija, Y. Tong, Z. Mi and S. Ouyang (2020). “Structural and functional insights into a novel two-component endolysin encoded by a single gene in Enterococcus faecalis phage.” PLoS Pathog 16(3): e1008394.

