## Supplementary Information for "Heterodimerization of Endolysin Isoforms During Bacterial Infection by Staphylococcal Phage φ2638A"

SUPPLEMENTARY TABLE S1. Oligonucleotides used for the amplification of genomic DNA fragments required for synthetic assembly of bacteriophage genomes.

| Fragment | Name | Primer sequence (5' → 3') | Phage | Template |
|---|---|---|---|---|
| f1 | 419 (F) | ACTATCGGAACGTCCAGATTTAGC | φ2638A $ply_{WT\text{-}HA}$ | φ2638A WT |
| | 412 (R) | TACCGACTGCGCGCATCTGAC | | |
| f2.1 | 413 (F) | TCTATCCGGAATGTAGCAGGTCAG | | |
| | HA_(R) | TATTTTAAGCGTAATCTGGAACATCGTATGGGTATTTAATTTCGCCCCACAACTTACCAACTTTAC | | |
| f3.1 | HA_(F) | ATTAAATACCCATACGATGTTCCAGATTACGCTTAAAATATGATATACTATGTATATCCACGACATGATTAG | | |
| | 416 (R) | CTTGATAGCCCAATGCCAATTCTG | | |
| f4 | 417 (F) | CAAAAAGGTAAAGTATCAGAATTGGCATTG | | |
| | 420 (R) | ATTGTTCAGGTCATACGCTAAATCTG | | |
| f2.2 | 413 (F) | TCTATCCGGAATGTAGCAGGTCAG | φ2638A $ply_{FL}$ | φ2638A WT |
| | CTC_(R) | GTTTGAATAGATATGTTTGAGCTCTTTCACGCTCCC | | |
| f3.2 | CTC_(F) | GAATGGGAGCGTGAAAGAGCTCAAACATATCTATTC | | |
| | 416 (R) | CTTGATAGCCCAATGCCAATTCTG | | |
| f3.3 | CTC_(F) | GAATGGGAGCGTGAAAGAGCTCAAACATATCTATTC | φ2638A $ply_{FL\text{-}HA}$ | φ2638A *ply-ha* |
| | 416 (R) | CTTGATAGCCCAATGCCAATTCTG | | |
| f2.4 | 413 (F) | TCTATCCGGAATGTAGCAGGTCAG | φ2638A $ply_{SV}$ | φ2638A WT |
| | Δ_M23_(R) | TTTGAATAGATATGTTTCATGTCACTTCAGCCCTTTCTCTTTAAGGTATTCC | | |
| f3.4 | Δ_M23_(F) | AGAAAGGGCTGAAGTGACATGAAACATATCTATTCAAACCATATTAAAGG | | |
| | 416 (R) | CTTGATAGCCCAATGCCAATTCTG | | |
| f2.5 | 413 (F) | TCTATCCGGAATGTAGCAGGTCAG | φ2638A $ply_{SV\text{-}HA}$ | φ2638A $ply_{WT\text{-}HA}$ |
| | Δ_M23_(R) | TTTGAATAGATATGTTTCATGTCACTTCAGCCCTTTCTCTTTAAGGTATTCC | | |
| f3.5 | Δ_M23_(F) | AGAAAGGGCTGAAGTGACATGAAACATATCTATTCAAACCATATTAAAGG | | |
| | 416 (R) | CTTGATAGCCCAATGCCAATTCTG | | |

SUPPLEMENTARY TABLE S2. Oligonucleotides and final protein expression plasmids

| Primer name and sequence (5'→3') | Template | Construct | Vector | 6×His[1)] |
|---|---|---|---|---|
| NdeI_Ply2638A_F<br>GATCCATATGCTAACTGCTATTGAC<br>Ply2638 _BamHI_R<br>TAATGGATCCTTATTTAATTTCGCCC | Ply2638A 1–486 | $Ply_{WT}$ | pET302 | - |
| NdeI_Ply2638A_F<br>GATCCATATGCTAACTGCTATTGAC<br>2638a_CTC_180_mut_F<br>GTGAAAGAGCTCAAACATATCTATTC<br>2638a_CTC_180_mut_R<br>GATATGTTTGAGCTCTTTCACGCTCC<br>Ply2638 _BamHI_R<br>TAATGGATCCTTATTTAATTTCGCCC | $Ply_{WT}$ | $Ply_{FL}$ | pET302 | - |
| Ply2638_short_NdeI_F<br>TCACCATATGAAACATATCTATTCAAACC<br>Ply2638 _BamHI_R<br>TAATGGATCCTTATTTAATTTCGCCC | $Ply_{WT}$ | $Ply_{SV}$ | pET302 | - |
| M23-2638_XhoI_F<br>ATAGCTCGAGATGCTGACC<br>M23-2638_BamHI_R<br>ATCGGGATCCTTAGTTTTTGC | Synthetic DNA[4)] | M23 | pET302 | NT |
| Ami(GA)_NdeI_F<br>TAATCACATATGGGTAGCGTGAAAGAGC<br>Ami(GA)_BamHI_R<br>AATTGGATCCTTAACCGTCGTAGTAATG TTTG | Synthetic DNA[4)] | Ami | pET302 | - |
| CBD2638_NdeI_F<br>ATAATACATATGTGGAAACAGAACAAAG ATGGC<br>CBD(GA2)_BamHI_R<br>ATTAGGATCCTTATTTGATTTCACCCCA CAG | Synthetic DNA[4)] | CBD | pET200 | - |
| - | - | HGFP[2)] | pQE30 | NT |
| - | - | HGFP_SH3b2638A[3)] | pQE30 | NT |
| - | - | HGFP_SH3bLST[3)] | pQE30 | NT |

[1)] Presence of the 6×Histidine tag: '-' no tag; 'NT' N-terminal His-tag

[2)] Loessner et al. (2002)

[3)] Doctoral Thesis, Fritz Eichenseher 2011, ETH Zurich

[4)] GeneArt (ThermoFisher)

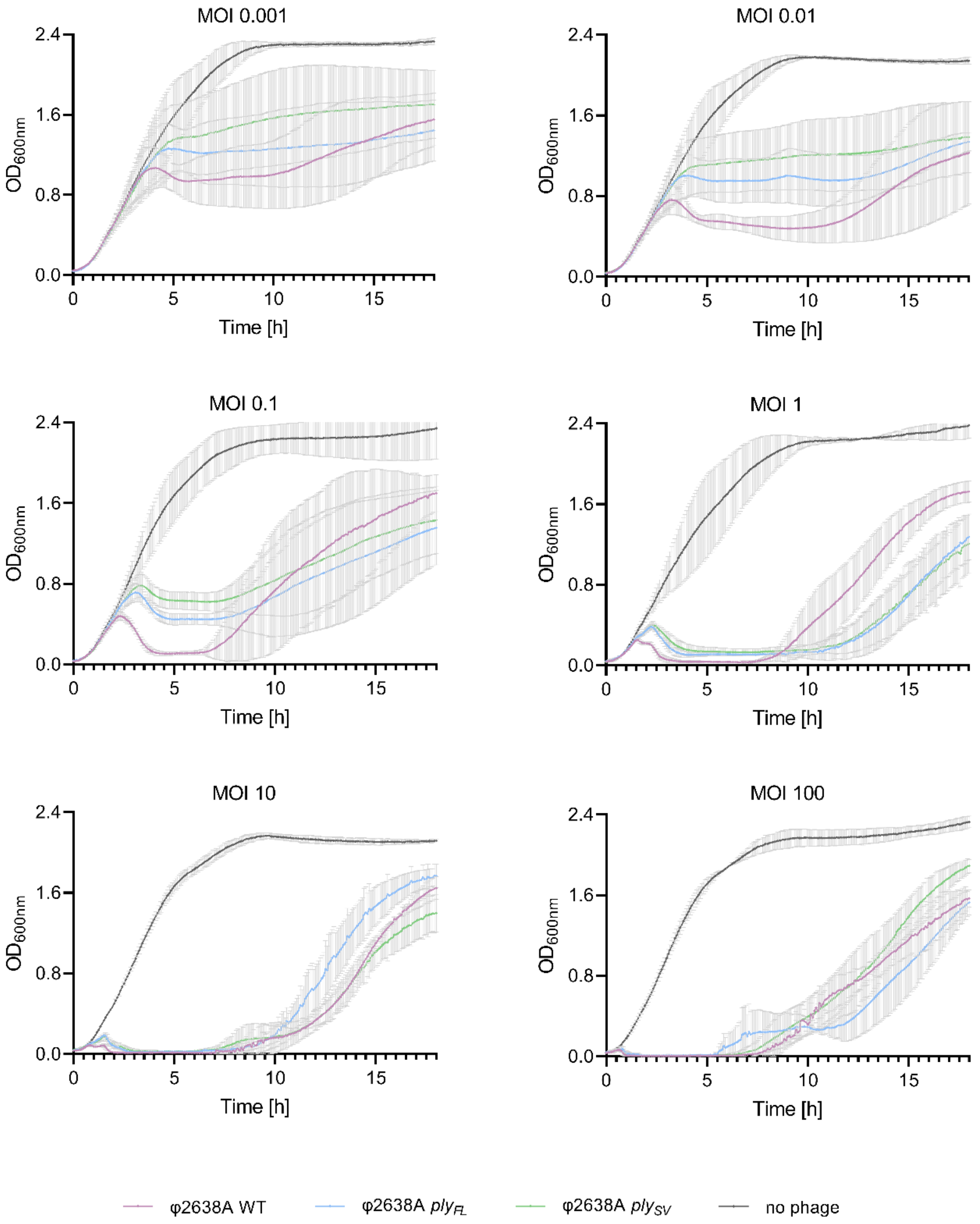


SUPPLEMENTARY FIGURE S1. **Turbidity reduction assays (TRAs) of wildtype and engineered φ2638A phages at different MOIs.** Wildtype φ2638A (purple), φ2638A*ply*$_{FL}$ (blue), φ2638A*ply*$_{SV}$ (green), or buffer alone as control (black) were added to $10^8$ CFU/mL *S. pseudointermedius* 2854 cells at different MOIs with bacteriolytic activity measured over 18 hours via optical density at $OD_{600nm}$. MOIs ranged from 0.001 to 100, corresponding to $10^5$ to $10^{10}$ PFU/mL of phages. The initial 8 hours of infection for the MOI 0.1 ($10^7$ PFU/mL) data are presented in FIGURE 1C.

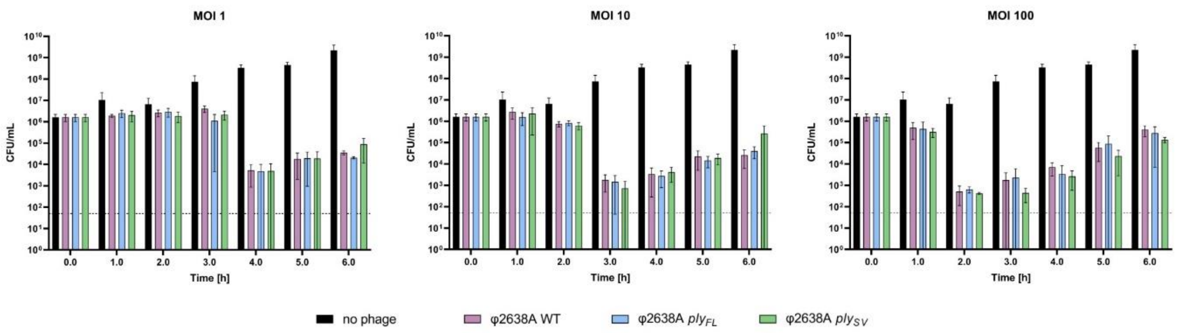


SUPPLEMENTARY FIGURE S2. **Time-kill assays (TKAs) of wildtype and engineered φ2638A phages at different MOIs.** Wildtype and engineered phages were added to 4 x $10^7$ CFU/mL *S. pseudointermedius* 2854 cells at different MOIs with bacteriolytic activity measured for six hours. Samples were taken, serially diluted, and plated on agar plates every hour. Surviving colonies were quantified after 16-hour incubation at 37 °C. MOIs ranged from 1.0 to 100, corresponding to $10^7$ to $10^{10}$ PFU/mL of phages.

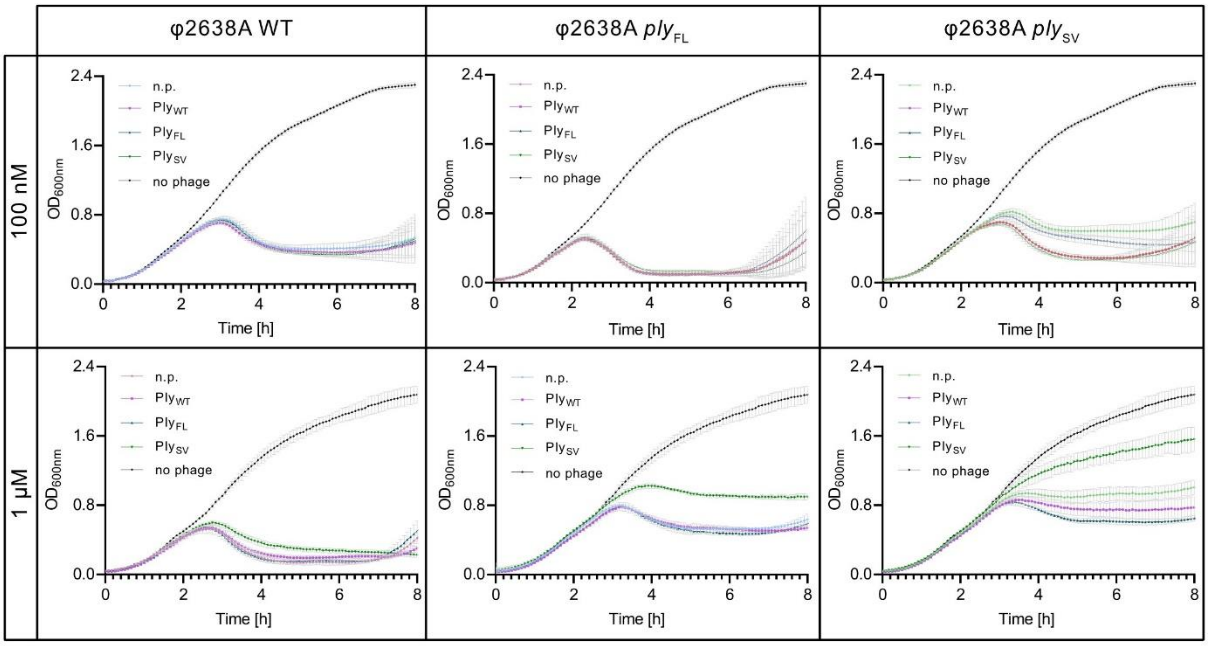


SUPPLEMENTARY FIGURE S3. **Turbidity reduction assays (TRAs) of wildtype and engineered φ2638A phages complemented with high amounts of recombinant endolysin**. Wildtype φ2638A and engineered phages, φ2638A*$ply_{FL}$* and φ2638A*$ply_{SV}$* were added to $10^8$ CFU/mL *S. pseudointermedius* 2854 cells at an MOI of 0.1 ($10^7$ PFU/mL) and complemented with 100 nM or 1 µM of recombinant $Ply_{WT}$ (the native mix of isoforms), $Ply_{FL}$, or $Ply_{SV}$, or no protein (n.p.) as control. Each experiment contains an uninfected bacterial culture (no phage) as negative control. Experiments with a 100 nM protein supplemented were performed in biological triplicates with technical triplicates. For TRAs with 1 µM protein supplemented only technical replicates were performed. Error bars represent the mean ± standard deviation.

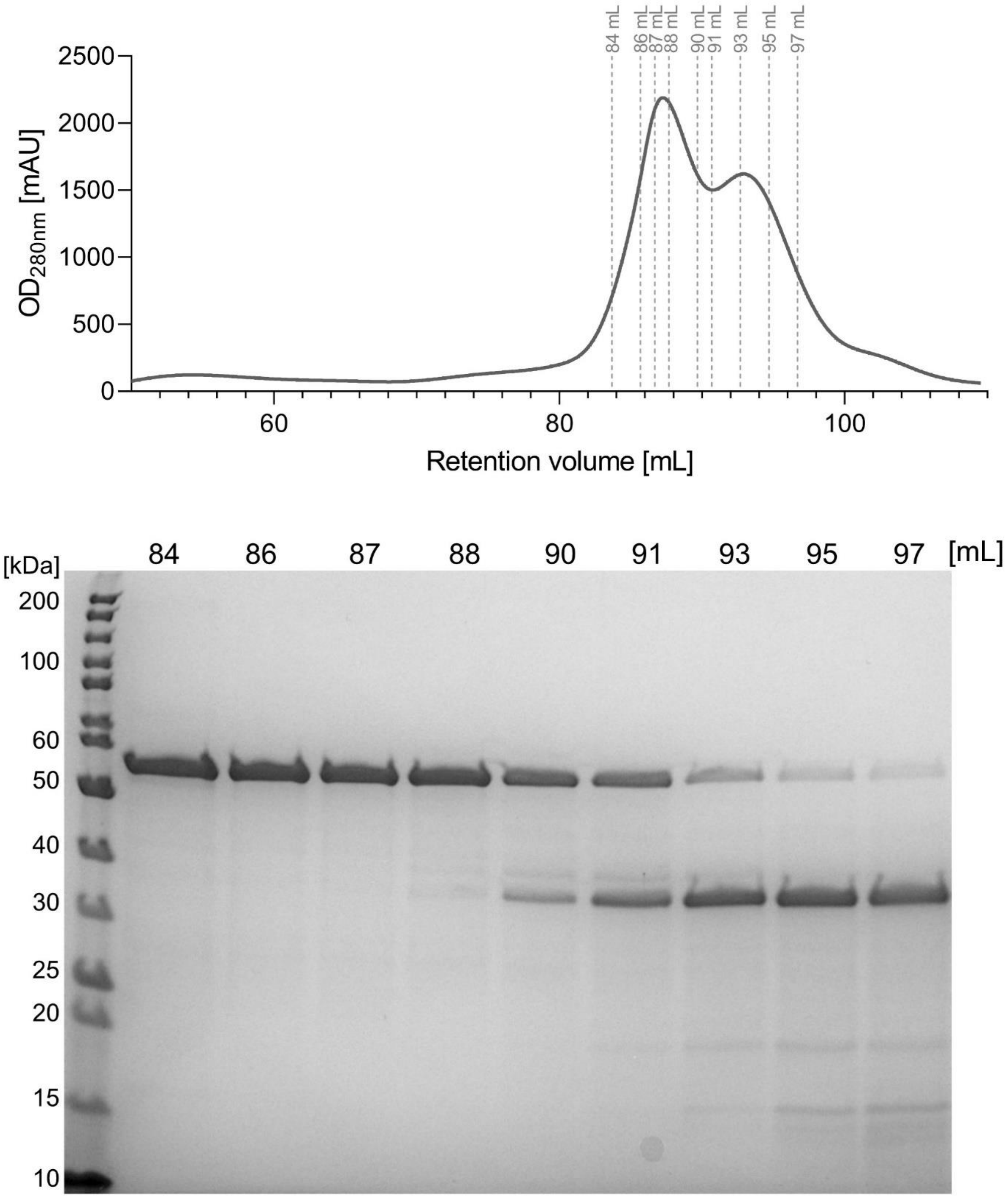


SUPPLEMENTARY FIGURE S4. **Size exclusion chromatography (SEC) Ply$_{WT}$ and SDS-PAGE gel of selected fractions.** Ply$_{WT}$ was purified via SEC leading to the observation of co-elution of FL and the SV isoform. 1 mL fractions were collected during the elution process and selected fractions indicated as dotted lines in the chromatogram were run on an SDS-PAGE gel.

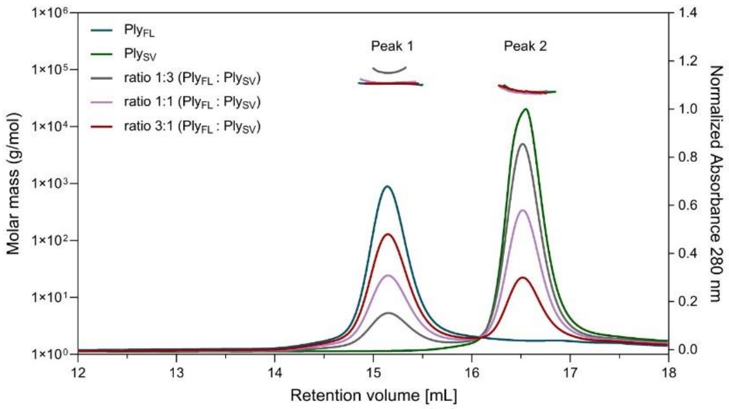


| Sample | Peak 1 | Peak 2 |
|---|---|---|
| Ply$_{FL}$ (55.5 kDa) | 56.4 (± 0.6) | - |
| Ply$_{SV}$ (35.3 kDa) | - | 40.8 (± 0.3) |
| Ply$_{FL}$ : Ply$_{SV}$ (1:3) | 97.0 (± 1.2) | 42.6 (± 0.4) |
| Ply$_{FL}$ : Ply$_{SV}$ (1:1) | 57.4 (± 0.5) | 42.5 (± 0.9) |
| Ply$_{FL}$ : Ply$_{SV}$ (3:1) | 59.1 (± 0.7) | 39.7 (± 0.4) |

SUPPLEMENTARY FIGURE S5. **Size exclusion chromatography with multi-angle static light scattering (SEC-MALS) analysis of Ply2638A isoform ratios.** The oligomeric states of Ply$_{FL}$, Ply$_{SV}$, and combinations of different ratios, 1:1, 3:1, 1:3 (w/w) were determined by SEC-MALS at a concentration of 1 mg/mL. The curves represent UV absorption at 280 nm, with the calculated molar mass (*y*-axis) for the respective peaks indicated above the corresponding peaks and summarized in the table on the right. BSA (bovine serum albumin) served as a reference standard during calibration.

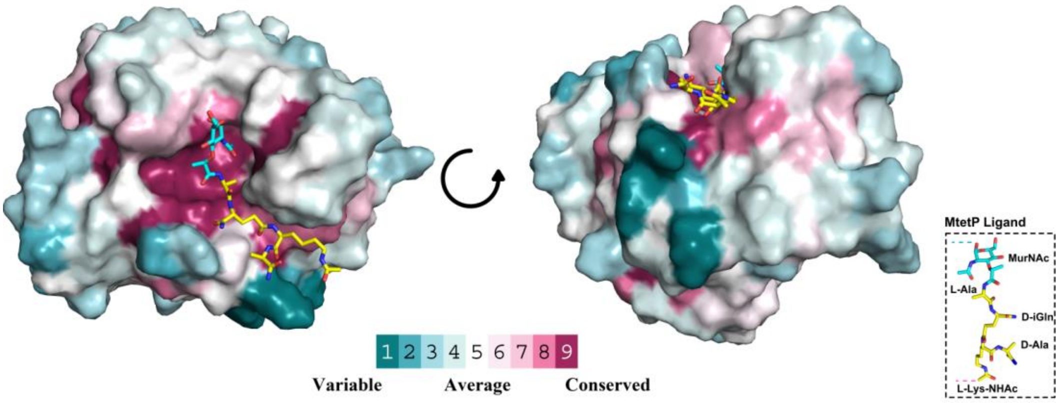


SUPPLEMENTARY FIGURE S6. **Amino acid conservation of the Ply2638A amidase domain.** Analysis was performed using the ConSurf server (Ashkenazy et al. 2016) with the default parameters for homolog search and multiple sequence alignment. Residues are colored according to the ConSurf conservation score (see legend). The muramyltetrapeptide (MtetP) ligand representative of *S. aureus* peptidoglycan was modelled into the active site by superpositioning with the MtetP co-crystallized structure of *S. aureus* autolysin, AmiA (PDB ID: 4KNL; Z-score 18.2; RMSD 2.3 Å) (Büttner et al. 2014).

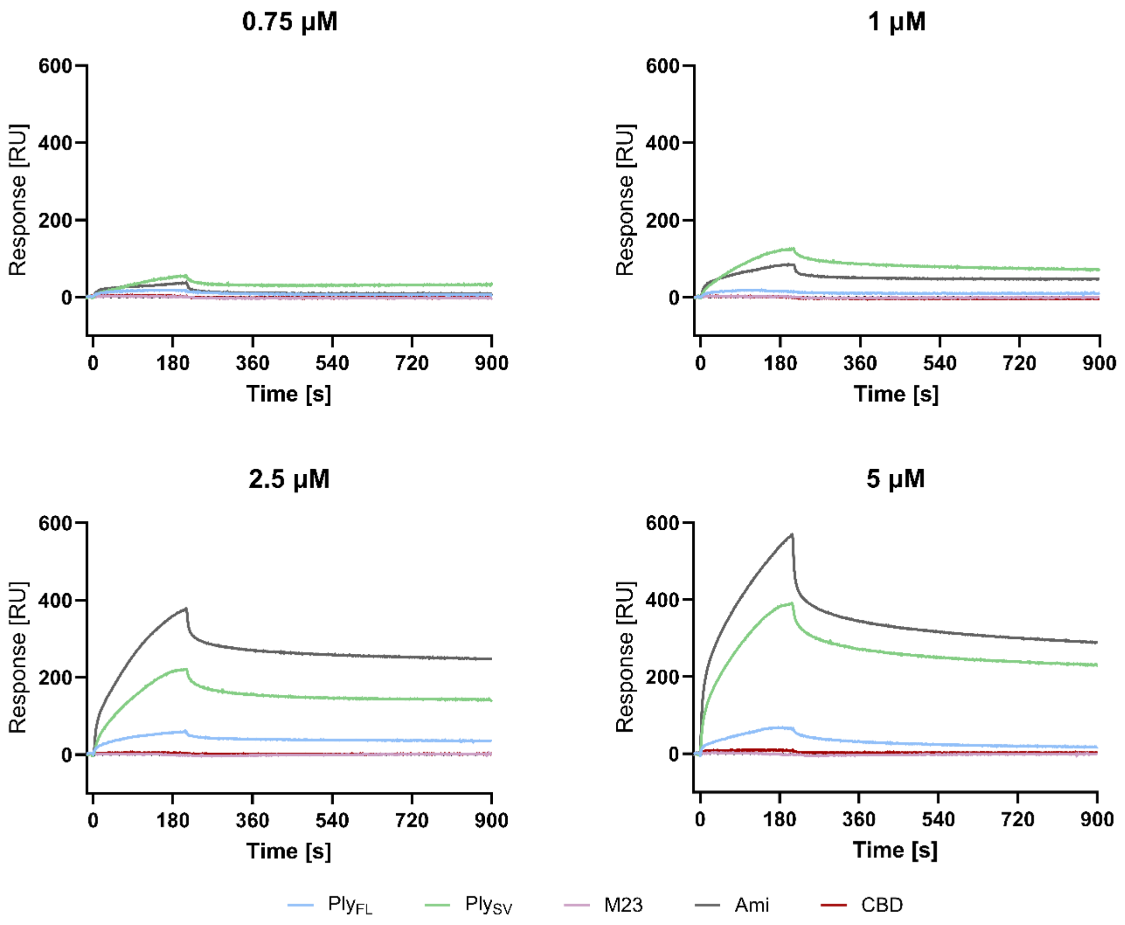


SUPPLEMENTARY FIGURE S7. **SPR data indicating interactions between amidase-containing constructs.** SPR sensorgrams of the analytes Ply$_{FL}$ (light blue), Ply$_{SV}$ (light green) and single domains M23 (purple), Ami (dark grey) and CBD (red) interacting with the ligand Ply$_{FL}$ immobilized on the chip surface. Four different analyte concentration were tested for all five constructs: 0.75 µM, 1 µM, 2.5 µM (shown in FIGURE 2E), and 5 µM.
